# Individualized multi-omic pathway deviation scores using multiple factor analysis

**DOI:** 10.1101/827022

**Authors:** Andrea Rau, Regina Manansala, Michael J. Flister, Hallgeir Rui, Florence Jaffrézic, Denis Laloë, Paul L. Auer

## Abstract

Malignant progression of normal tissue is typically driven by complex networks of somatic changes, including genetic mutations, copy number aberrations, epigenetic changes, and transcriptional reprogramming. To delineate aberrant multi-omic tumor features that correlate with clinical outcomes, we present a novel pathway-centric tool based on the multiple factor analysis framework called *padma*. Using a multi-omic consensus representation, *padma* quantifies and characterizes individualized pathway-specific multi-omic deviations and their underlying drivers, with respect to the sampled population. We demonstrate the utility of *padma* to correlate patient outcomes with complex genetic, epigenetic, and transcriptomic perturbations in clinically actionable pathways in breast and lung cancer.

## 1. Introduction

Large sets of patient-matched multi-omics data have become widely available for large-scale human health studies in recent years, with notable examples including the The Cancer Genome Atlas (TCGA; The Cancer Genome Atlas Research Network *and others*., 2013) and Trans-omics for Precision Medicine (TOPMed) program. The increasing emergence of multi-omic data has in turn led to a renewed interest in multivariate, multi-table approaches (Meng *and others*, 2016) to account for interdependencies within and across data types (Husson *and others*, 2017). In such large-scale multi-level data, there is often limited or incomplete *a priori* knowledge of relevant phenotype groups for comparisons, and a primary goal may be to identify subsets of individuals that share common molecular characteristics, design therapies in the context of personalized medicine, or identify relevant biological pathways for follow-up. With these goals in mind, many multivariate approaches have the advantage of being unsupervised, using matched or partially matched omics data across genes, obviating the need for predefined groups for comparison as in the framework of standard differential analyses. A variety of such approaches has been proposed in recent years. For example, *Multi-omics Factor Analysis* (MOFA) uses group factor analysis to infer sets of hidden factors that capture biological and technical variability for downstream use in sample clustering, data imputation, and sample outlier detection (Argelaguet *and others*, 2018).

In multi-omic integrative analyses, an intuitive first approach is to consider a gene-centric analysis, as we previously proposed in the *EDGE in TCGA* tool (Rau *and others*, 2018). Expanding such analyses to the pathway-level is also of great interest, as it can lead to improved biological interpretability as well as reduced or condensed gene lists to facilitate the generation of relevant hypotheses. In particular, our goal is to define a method that quantifies an individual’s deviation from a sample average, at the pathway-level, while simultaneously accounting for multiple layers of molecular information. Several related approaches for pathway-specific single-sample analyses have been proposed in recent years (Vaske *and others*, 2010; Verbeke *and others*, 2015; Drier *and others*, 2013). For example, *PARADIGM* (Vaske *and others*, 2010) is a widely used approach based on structured probabilistic factor graphs to prioritize relevant pathways involved in cancer progression as well as identify patient-specific alterations; both pathway structures and multi-omic relationships are hard-coded directly in the model, but it requires a discretization of the data and is now a closed-source software, making extensions and application to other gene sets difficult. *Pathway relevance ranking* (Verbeke *and others*, 2015) integrates binarized tumor-related omics data into a comprehensive network representation of genes, patient samples, and prior knowledge to calculate the relevance of a given pathway to a set of individuals. A pathway-centric supervised principal component-based analysis implemented in *pathwayPCA* (Odom *and others*, 2019) performs gene selection and estimates latent variables for association testing with respect to binary, continuous, and survival outcomes within each set of omics data independently. *Pathifier* (Drier *and others*, 2013) instead seeks to calculate a personal pathway deregulation score (PDS), based on the distance of a single individual from the median reference sample on a principal curve; this principal curve approach is analogous to a nonlinear principal components analysis (PCA), but can be applied only to a single-omic dataset (e.g., gene expression). For both *PARADIGM* and *Pathifier*, clusters of scores across pathways are shown to correlate with a clinically relevant clustering of patients.

Here, we extend the basic philosophy of the *Pathifier* approach to multi-omics data, using an innovative application of a Multiple Factor Analysis (MFA), to quantify individualized pathway deviation scores. In particular, we propose an approach called *padma* (“PAthway Deviation scores using Multiple factor Analysis”) to characterize individuals with aberrant multi-omic profiles for a given pathway of interest and to quantify this deviation with respect to the sampled population using a multi-omic consensus representation. We further investigate the following succession of questions. In which pathways are high deviation scores strongly associated with measures of poor prognosis? For such pathways, which specific individuals are characterized by the most highly aberrant multi-omic profile? And for such individuals, which specific genes and omics drive large pathway deviation scores? By providing graphical and numerical outputs to address these questions, *padma* represents both an approach for generating hypotheses as well as an exploratory data analysis tool for identifying individuals and genes/omics of potential interest for a given pathway.

There is already some precedent for using MFA to integrate multi-omic data, although existing approaches differ from that proposed here. For instance, de Tayrac *and others* (2009) suggested using MFA for paired CGH array and microarray data, superimposed with functional gene ontology terms, to highlight common structures and provide graphical outputs to better understand the relationships between omics. In addition, *padma* shares some similarities with a recently proposed integrative multi-omics unsupervised gene set analysis called *mogsa*, which is similarly based on a MFA (Meng *and others*, 2019). By calculating an integrated multi-omics enrichment score for a given gene set with respect to the full gene list, *mogsa* identifies gene sets driven by features that explain a large proportion of the global correlated information among different omics. In addition, these integrated enrichment scores can be decomposed by omic and used to identify differentially expressed gene sets or reveal biological pathways with correlated profiles across multiple complex data sets. However, the fundamental difference in the two approaches is that *mogsa* evaluates pathway-specific enrichment with respect to the entire set of genes, while *padma* instead focuses on identifying and quantifying pathway-specific multi-omic deviations between each individual and the sampled population.

## 2. Methods

### 2.1 Pathway-centric multiple factor analysis for multi-omic data

MFA represents an extension of principal component analysis for the case where multiple quantitative data tables are to be simultaneously analyzed (Escofier and Pagès, 2014; Pagès, 2015; Lê *and others*, 2008; Abdi *and others*, 2013). As such, MFA is a dimension reduction method that decomposes the set of features from a given gene set into a lower dimension space. In particular, the MFA approach weights each table individually to ensure that tables with more features or those on a different scale do not dominate the analysis; all features within a given table are given the same weight. These weights are chosen such that the first eigenvalue of a PCA performed on each weighted table is equal to 1, ensuring that all tables play an equal role in the global multi-table analysis. According to the desired focus of the analysis, data can be structured either with molecular assays (e.g., RNA-seq, methylation, miRNA-seq, copy number alterations) as tables (and genes as features within omics), or with genes as tables (and molecular assays as features within genes). The MFA weights balance the contributions of each omic or of each gene, respectively. In this work, we focus on the latter strategy in order to allow different omics to contribute to a varying degree depending on the chosen pathway. In addition, we note that because the MFA is performed on standardized features, simple differences in scale between omics (e.g., RNA-seq log-normalized counts versus methylation logit-transformed beta values) do not impact the analysis.

More precisely, consider a pathway or gene set composed of *p* genes (Figure 1A), each of which is measured using up to *k* molecular assays (e.g., RNA-seq, methylation, miRNA-seq, copy number alterations), contained in the set of gene-specific matrices *X*_1_,…, *X_p_* that have the same *n* matched individuals (rows) and *j*_1_,…, *j_p_* potentially unmatched variables (columns) in each, where *j_g_* ∈ {1,…, *k*} for each gene *g* = 1,…, *p*. Because only the observations and not the variables are matched across data tables, genes may be represented by potentially different subset of omics data (e.g., only expression data for one gene, and expression and methylation data for another).

**Fig. 1.**
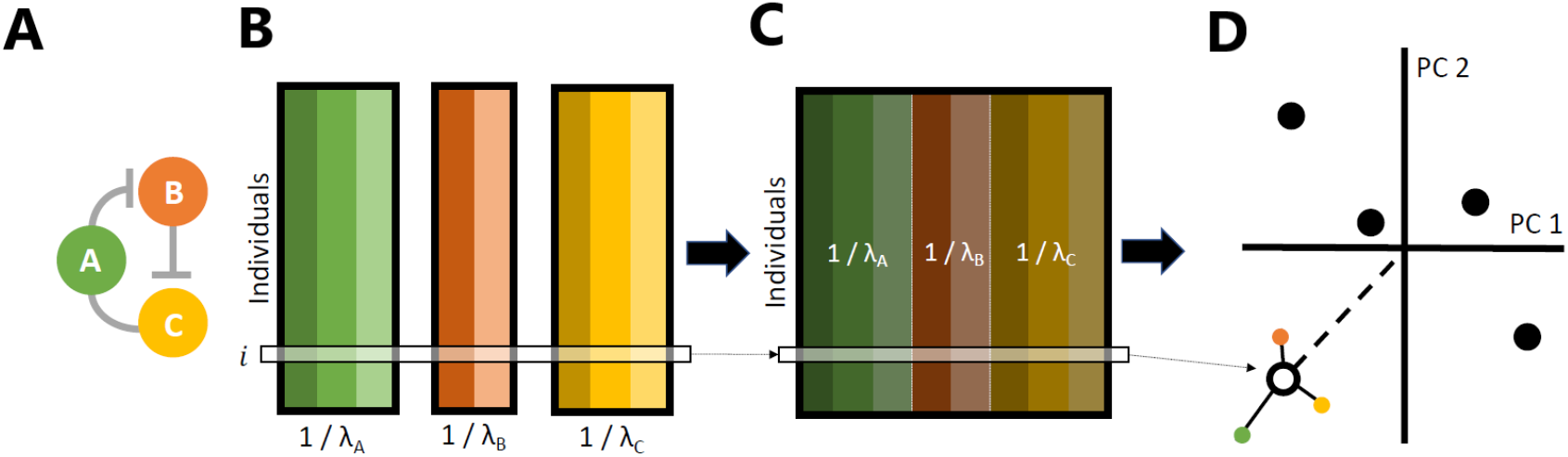
Illustration of the padma approach for calculating individualized multi-omic pathway deviation scores. (A-B) For a given pathway, matched multi-omic measures for each gene are assembled, with individuals in rows. Note that genes may be assayed for varying types of data (e.g., measurements for one gene may be available for expression, methylation, and copy number alterations, while another may only have measurements available for expression and methylation). (C) Using a Multiple Factor Analysis, each gene table is weighted by its largest singular value, and per-gene weighted tables are combined into a global table, which in turn is analyzed using a Principal Component Analysis. (D) Finally, each individual *i* is projected onto the consensus pathway representation; the individualized pathway deviation score is then quantified as the distance of this individual from the average individual. These scores can be further decomposed into parts attributed to each gene in the pathway.

In the first step, these data tables are generally standardized (i.e., centered and scaled). Next, an individual PCA is performed using singular value decomposition for each gene table *X_g_*, and its largest singular value 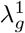, which corresponds to the variance of the first principal component, is calculated (Figure 1B). Note that 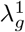 represents a function of both the number of variables in a given table and the redundancy among them; the more redundant a set of variables, the less new information is contributed by each given the others, and the larger 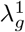 will be. Then, all features in each gene table *X_g_* are weighted by 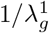, and a global PCA is performed using a singular value decomposition on the concatenated set of weighted standardized tables, 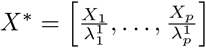 (Figure 1C). Using this weighting scheme, genes with highly correlated measures among some combination of multi-omic assays will thus tend to be down-weighted in the global analysis, while those with complementary information contributed by different assays will tend to be upweighted. Although this is the standard weighting used in MFA, we note that other potential strategies do exist, for example in the consensus PCA (Wold *and others*, 1987), which uses as weights the inverse of the total inertia of each table (equal to the number of variables in the case of standardized variables).

The global PCA performed on the weighted standardized data yields a matrix of components (i.e., latent variables) in the observation and variable space. Optionally, an independent set of supplementary individuals (or supplementary variables) can then be projected onto the original representation; this is performed by centering and scaling variables for the supplementary individuals (or individuals for the supplementary variables, respectively) to the same scale as for the reference individuals, and projecting these rescaled variables into the reference PCA space. Note that in the related *mogsa* approach, supplementary binary variables representing gene membership in gene sets are projected onto a transcriptome-wide multiple factor analysis to calculate gene set scores (Meng *and others*, 2019).

The MFA thus provides a consensus across-gene representation of the individuals for a given pathway, and the global PCA performed on the weighted gene tables decomposes the consensus variance into orthogonal variables (i.e., principal components) that are ordered by the proportion of variance explained by each. The coordinates of each individual on these components, also referred to as factor scores, can be used to produce factor maps to represent individuals in this consensus space such that smaller distances reflect greater similarities among individuals. In addition, partial factor scores, which represent the position of individuals in the consensus for a given gene, can also be represented in the consensus factor map; the average of partial factor scores across all dimensions and genes for a given individual corresponds to the factor score (Figure 1D). A more thorough discussion of the MFA, as well as its relationship to a PCA and additional details about the calculation of factor scores and partial factor scores, may be found in the Supplementary Methods.

### 2.2 Individualized pathway deviation scores

In the consensus space obtained from the MFA, the origin represents the “average” pathway behavior across genes, omics, and individuals; individuals that are projected to increasingly distant points in the factor map represent those with increasingly aberrant values, with respect to this average, for one or more of the omics measures for one or more genes in the pathway. To quantify these aberrant individuals, we propose an individualized pathway deviation score *d_i_* based on the multidimensional Euclidean distance of the MFA component loadings for each individual to the origin:

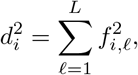

where *f_i,ℓ_* corresponds to the MFA factor score of individual *i* in component *ℓ*, and *L* corresponds to the rank of *X*\*. Note that this corresponds to the weighted Euclidean distance of the scaled multi-omic data (for the genes in a given pathway) of each individual to the origin. These individualized pathway deviation scores are thus nonnegative, where smaller values represent individuals for whom the average multi-omic pathway variation is close to the average, while larger scores represent individuals with increasingly aberrant multi-omic pathway variation with respect to the average. An individual with a large pathway deviation score is thus characterized by one or more genes, with one or more omic measures, that explain a large proportion of the global correlated information across the full pathway.

Note that the full set of components is used for this deviation calculation, rather than subsetting to an optimal number of components; we remark that due to their small variance relative to lower components, higher components contribute relatively little to the overall pathway deviation scores. When all components are used in *padma*, there is no dimension reduction and the calculation of the pathway deviation score (and the corresponding gene-level contributions) is equivalent to that calculated in the original space of the concatenated set of weighted, standardized variables. However, if desired, the user can calculate the pathway deviation score on a subset of components, for example after removing one or more components that are correlated with batch effects or those that explain little variability. Finally, to facilitate comparisons of scores calculated for pathways of differing sizes (e.g., the number of genes), deviation scores with respect to the origin are normalized for the pathway size by dividing them by the number of genes in the pathway.

### 2.3 Decomposition of individualized pathway deviation scores into per-gene contributions

In order to quantify the role played by each gene for each individual, we decompose the individualized pathway deviation scores into gene-level contributions. Recall that the average of partial factor scores across all MFA dimensions corresponds to each individual’s factor score. We define the gene-level deviation for a given individual as follows:

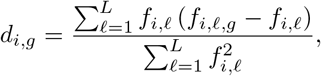

where as before *f_i,ℓ_* corresponds to the MFA factor score of individual *i* in component *ℓ, L* corresponds to the rank of *X*\*, and *f_i,ℓ,g_* corresponds to the MFA partial factor score of individual *i* in gene *g* in component *ℓ*. Note that by construction, the contributions of all pathway genes to the overall deviation score sum to 0. In particular, per-gene contributions can take on both negative and positive values according to the extent to which the gene influences the deviation of the overall pathway score from the origin (i.e., the global center of gravity across individuals); large positive values correspond to tables with a large influence on the overall deviation of an individual, while large negative values correspond to genes that tend to be most similar to the global average. In the following, we additionally scale these per-gene scores by the inverse overall pathway score to highlight genes with highly atypical multi-omic measures both with respect to other genes in the pathway and with respect to individuals in the population.

Interestingly, it is possible to quantify the contribution of any single variable of any arbitrary grouping of variables (e.g. individual omics, all assays for a given gene family) to the individualized pathway deviation score; see the Supplementary Methods for more details.

### 2.4 Quantifying percent contribution of omics to pathway-centric multiple factor analysis

The richness of MFA outputs also includes various decompositions of the total variance (that is, the sum of the variances of each individual MFA component) of the multi-omic data for a given pathway. Similarly to a standard PCA, the percent contribution of each axis of the MFA can be calculated as the ratio between the variance of the corresponding MFA component and the total variance; by construction, the fraction of explained variance explained decreases as the MFA dimension increases. Similarly, the percent contribution to the inertia of each axis for a given omic, gene, or individual can be quantified as the ratio between the inertia of its respective partial projection in the consensus space and the inertia of the full data projection for that axis. These per-gene, per-omic, and per-individual contributions can be quantified for a subset of components (e.g., the first ten dimensions) or for the entire set of components; here, as we calculate individualized pathway deviation scores using the full set of dimensions, we also calculated a weighted per-omic contribution, which corresponds to the average contribution across all dimensions, weighted by the corresponding eigenvalue.

### 2.5 padma R software package

The proposed method described above has been implemented in an open-source R package called *padma*, freely available at https://github.com/andreamrau/padma. *Padma* notably makes use *FactoMineR* (Lê *and others*, 2008; Husson *and others*, 2017) to run the MFA; heatmaps in the following results were produced using *ComplexHeatmap* (Gu *and others*, 2016). All of the analyses in this paper were performed using R v3.5.1. In addition, all R scripts used to generate the results in this work may be found at https://github.com/andreamrau/RMFRJLA_2019.

## 3. Application

### 3.1 Description of TCGA data and pathway collection

We illustrate the utility of *padma* on data from two cancer types with sufficiently large multi-omic sample sizes in the TCGA database: invasive breast carcinoma (BRCA), which was chosen as individuals have previously been classified (Paquet and Hallett, 2015) into one of five molecular subtypes (Luminal A, Luminal B, Her2+, Basal, and Normal-like), as well as lung adenocarcinoma (LUAD), which was chosen for its high recorded mortality. The multi-omic TCGA data were downloaded and processed as described in Rau *and others* (2018); in particular, all associated scripts can be found at https://doi.org/10.5281/zenodo.3524080 and additional details are provided in the Supplementary Methods. In this study, pre-processed and batch-corrected multi-omic data for BRCA and LUAD included gene expression, methylation, copy number alterations (CNA), and microRNA (miRNA) abundance for *n* = 504 and *n* = 144 individuals, respectively.

The *padma* approach integrates multi-omic data by mapping omics measures to genes in a given pathway. Although this assignment of values to genes is straightforward for RNA-seq, CNA, and methylation data, a definitive mapping of miRNA-to-gene relationships does not exist, as miRNAs can each potentially target multiple genes. Many methods and databases based on textmining or bioinformatics-driven approaches exist to predict miRNA-target pairs (Riffo-Campos *and others*, 2016). Here, we make use of the curated miR-target interaction (MTI) predictions in miRTarBase version 7.0 (Chou *and others*, 2018), using only exact matches for miRNA IDs and target gene symbols and predictions with the “Functional MTI” support type. Although the TCGA data used here have been filtered to include only those genes for which expression measurements are available, there are cases where missing values are recorded in other omics datasets (e.g., when no methylation probe was available in the promoter region of a gene, or when no predicted MTIs were identified) or where a given feature has little or no variance across individuals. In this analysis, features for a given omics dataset were removed from the analysis only if missing values are recorded for all individuals or if the feature has minimal variance across all individuals (defined here as < 10 ^−5^ before scaling); any remaining missing values are mean-imputed, although more sophisticated imputation strategies, such as those proposed in the *missMDA* package, could be used instead (Josse and Husson, 2016). After running *padma*, we remark that the first ten MFA dimensions represent a modest proportion of the total multi-omic variance across pathways for both cancers (Supplementary Figure 5; BRCA median = 46.1%, LUAD median = 51.9%); the number of MFA components needed to explain 80% of the total variability was strongly associated with the total number of features in each pathway (Supplementary Figure 9).

As a measure of patient prognosis, we focused on two different metrics. First, we used the standardized and curated clinical data included in the TCGA Pan-Cancer Clinical Resource (Liu *and others*, 2018) to identify the progression-free interval (PFI). The PFI corresponds to the period from the date of diagnosis until the date of the first occurrence of a new tumor event (e.g., locoregional recurrence, distant metastasis) and typically has a shorter minimum follow-up time than measures such as overall survival. In the BRCA data, a total of 72 uncensored and 434 censored events were recorded (median PFI time of 792 and 915 days, respectively); among LUAD individuals, a total of 65 uncensored and 79 censored events were recorded (median PFI time of 439 and 683 days, respectively). Second, we downloaded the histological grade for breast cancer (http://legacy.dx.ai/tcga_breast on March 7, 2019), which is an established cancer hallmark of cellular de-differentiation and poor prognosis (Heng *and others*, 2017). Tumors are typically graded by pathologists on a scale of 1 (well-differentiated), 2 (moderately differentiated), or 3 (poorly differentiated) based on three different measures, including nuclear pleomorphism, glandular/tubule formation, and mitotic index, where higher grades correspond to faster-growing cancers that are more likely to spread (Supplementary Table 3).

Finally, we focus our attention on a collection of 1136 pathways included in the MSigDB canonical pathways curated gene set catalog (Liberzon *and others*, 2011), which includes genes whose products are involved in metabolic and signaling pathways reported in curated public databases; additional details on these pathways may be found in the Supplementary Methods.

### 3.2 Computational validation of pathway deviation scores

Before exploring in detail the results of *padma* on the TCGA breast and lung tumor samples, we first sought to computationally validate the extent to which the pathway deviation scores correctly identify observations known to have aberrant multi-omic profiles. Specifically, we made use of the 70 matched healthy tissue samples in the TCGA breast cancer data for which RNA-seq, miRNA-seq, and methylation assays were available (as copy number alterations are called by comparing tumor to healthy tissue, these are not available for healthy tissue). These multi-omic healthy tissue samples were subsequently batch-corrected in the same way as the tumor samples. Next, to create a multi-omic data set with a set of “true positives”, we randomly selected 5 tumor samples to include with the 70 healthy samples; for each pathway, this random sampling was repeated 20 independent times. Using the RNA-seq, miRNA-seq, and methylation data for the full set of 75 samples, we then evaluated whether the *padma* pathway deviation scores were able to successfully identify the 5 tumor samples by calculating the Area Under the Curve (AUC) of the Receiver Operating Characteristic (ROC) curve for each of the 1136 pathways. Mean AUC values across the 20 repetitions were very high for nearly all pathways considered (25% quantile = 0.988; median = 0.993; 75% quantile = 0.996; Supplementary Figure 7). As a whole, this suggests that the *padma* deviation scores indeed reflect true biological signal, as the tumor controls added to healthy samples nearly always had the largest deviation scores across pathways.

In addition, we conducted a simulation study to investigate the conditions (i.e., sample size, percentage of aberrant individuals in the population, and number of driver genes and omics) under which *padma* pathway deviation scores can correctly identify aberrant individuals. We compared the AUC values of *padma* with those of pathway deviation scores calculated using two alternatives: a PCA on all concatenated data tables (i.e., where MFA per-table weights were not applied) and a *padma* single-omics approach. Full details of the simulation study may be found in the Supplementary Methods, and results are shown in Supplementary Figures 12-14 and Supplementary Table 6. Overall, we confirm the solid performance of *padma* across a wide range of settings, as well as an advantage in identifying aberrant individuals for *padma* compared to a PCA-based alternative, particularly in cases with smaller sample sizes (i.e., n 50) and fewer driver genes and omics. Finally, we also confirmed that the per-gene contributions to the individualized *padma* pathway deviation scores successfully recover the true gene drivers in all scenarios considered.

### 3.3 Large deviation scores for relevant oncogenic pathways are associated with survival in lung cancer

The first major question we address is the prioritization of pathways that are associated with a given phenotype of interest. After processing the TCGA data and assembling the collection of gene sets, we sought to identify a subset of pathways for which deviation scores were significantly associated with patient outcome, as measured by PFI. To focus on pathways with the largest potential signal (i.e., those for which a small number of individuals have very large deviation scores relative to the remaining individuals) we consider only those with the most highly positively skewed distribution of deviation scores. For each of the top 5% of pathways (*n* = 57) ranked according to their Pearson’s moment coefficient of skewness, we fit a Cox proportional hazards (PH) model for the PFI on the pathway deviation score, additionally controlling for age at initial pathologic diagnosis (minimum = 42; median = 68; maximum = 86), gender (88 females, 56 males), and American Joint Committee on Cancer (AJCC) pathologic tumor stage (Stage I, n = 80; Stage II, n = 33; Stage III+, n = 31). Using the Benjamini-Hochberg (BH; Benjamini and Hochberg, 1995) adjusted p-values from a likelihood ratio test (FDR < 5%), we identified 32 pathways with deviation scores that were significantly associated with the progression-free interval in lung cancer (Supplementary Table 1; see Supplementary Table 4 for the full gene lists in each pathway); for all of these, higher pathway scores corresponded to a worse survival outcome. Although overlaps in gene lists between pairs of pathways create a positive dependency structure among the deviation scores (Supplementary Figure 11), the BH correction method has been shown to control the FDR in such families of tests (Benjamini and Yekutieli, 2001). Note that the filtering on skewness of the pathway scores is performed completely independently of the survival phenotype, ensuring that the downstream survival analysis is not biased (Bourgon *and others*, 2010). Of note, while candidates within the majority of deviated pathways (Supplementary Table 1) have been univariately associated with patient outcome (e.g., cell cycle, DNA repair, and apoptosis; Bosken *and others*, 2002; Singhal *and others*, 2005), the *padma* TCGA analysis is unique in its ability to extend these associations across multiple gene patient-specific perturbations within a pathway at the genomic and transcriptomic RNA levels.

It is also of interest to evaluate the difference in results provided by a multi-omic versus single-omic pathway deviation analysis. To this end, we used *padma* to calculate deviation scores and fit the corresponding Cox PH survival analysis for the same n=57 pathways using RNA-seq lung cancer data alone (note that this corresponds to running an MFA with a single omic feature in each gene table). Pathway deviation scores for each individual were moderately correlated between the single- and multi-omic analyses (Pearson correlation; minimum = 0.56, median = 0.67, max = 0.82), but the majority of the pathways had considerably smaller p-values in the multi-omic analysis as compared to the single-omic analysis (Supplementary Figure 8); further, after BH correction none of the 57 pathways were significantly associated with survival (FDR < 5%) in the single-omic analysis. Although single-omic deviation scores for pathways other than those studied here may be significantly associated with survival, this result does indicate that a multivariate analysis of multi-omic data does capture a different signal than does a single-omic analysis.

The detection of several pathways related to DNA repair (ATM, Homologous DNA repair, BRCA1/2-ATR; Supplementary Table 1), as well as cell cycle and apoptosis related pathways, prompted us to consider whether these pathway deviation scores are simply acting as proxies for the tumor mutational burden (i.e., the total number of nonsynonymous mutations) for each individual. To investigate this, we estimated the mutational burden for each individual by counting the number of somatic nonsynonymous mutations in a set of cancer-specific driver genes (*n* = 183 and n = 181 genes in breast and lung cancer, respectively) identified by IntOGen (Gonzalez-Perez *and others*, 2013). After adding a constant of 1 to these counts and log-transforming them, we fit a linear model to evaluate their association with the pathway deviation scores; after correcting p-values from the Wald test statistic for multiple testing (FDR < 10%), no pathways were found to be associated with the mutational burden. In addition, when repeating the Cox PH model described above including the log-mutational burden as an additional covariate, raw p-values were generally similar to previous values (Spearman correlation: *ρ* = 0.9981). This suggests that the biological signal contained in the pathway deviation scores is indeed independent of that linked to mutational burden.

### 3.4 Padma identifies individualized aberrations in the Df-GDP dissociation inhibitor signaling pathway in lung cancer

To illustrate the full range of results provided by *padma*, we focus in particular on the results for the D4-GDP dissociation inhibitor (GDI) signaling pathway. D4-GDI is a negative regulator of the ras-related Rho Family of GTPases, and it has been suggested that it may promote breast cancer cell proliferation and invasiveness (Zhang *and others*, 2009; Zhang and Zhang, 2006). The D4-GDI signaling pathway is made up of 13 genes; RNA-seq, methylation, and CNA measures are available for all 13 genes, with the exception of CYCS and PARP1, for which no methylation probes were measured the promoter region. In addition, miRNA-seq data were included for one predicted target pair: hsa-mir-421 → CASP3. Over the 13 genes in the pathway, 130 of the 144 individuals had no nonsynonymous mutations, while 13 and 1 individuals had 1 or 3 such mutations; ARHGAP5 and CASP3 were most often characterized by mutations (3 individuals affected for each). Notably, although the D4-GDI pathway has been previously implicated in breast cancer aggressiveness (Zhang *and others*, 2009; Zhang and Zhang, 2006), this is to our knowledge the first evidence suggesting that D4-GDI pathway might play a similar role in promoting lung cancer.

Using the multi-omic data available for the D4-GDI signaling pathway, we can use the outputs of *padma* to better understand the individualized drivers of multi-omic variation. In particular, it is possible to quantify both gene-specific deviation scores as well as an overall pathway deviation score for each individual, respectively based on the set of partial or full MFA components. We first visualize the scaled gene-specific deviation scores for the top and bottom decile of individuals, according to their overall pathway deviation score (Figure 2); these groups thus correspond to the individuals that are least and most similar to the average individual within the population. We remark that the 10% of individuals with the most aberrant overall scores for the D4-GDI signaling pathway, who also had a high 1- and 5-year mortality rate, are those that also tend to have large aberrant (i.e., red in the heatmap) scaled gene-specific deviation scores for one or more genes. For example, the two individuals with the largest overall scores, TCGA-78-7536 and TCGA-78-7155 (12.79 and 12.31, respectively), both had large scaled gene-specific scores for CASP3 (12.93 and 17.05, respectively), CASP1 (27.80 and 10.85, respectively), and CASP8 (29.72 and 22.61, respectively). While a subset of five individuals from the top decile were all characterized by high deviation scores for JUN (TCGA-64-5775, TCGA-55-6972, TCGA-50-5051, TCGA-44-6779, TCGA-49-4488), several other genes appear to have relatively small deviation scores for all individuals plotted here (e.g., PRF1, PARP1). In addition, we remark the presence of highly individualized gene-specific aberrations (e.g., APAF1 in individual TCGA-55-7725).

**Fig. 2.**
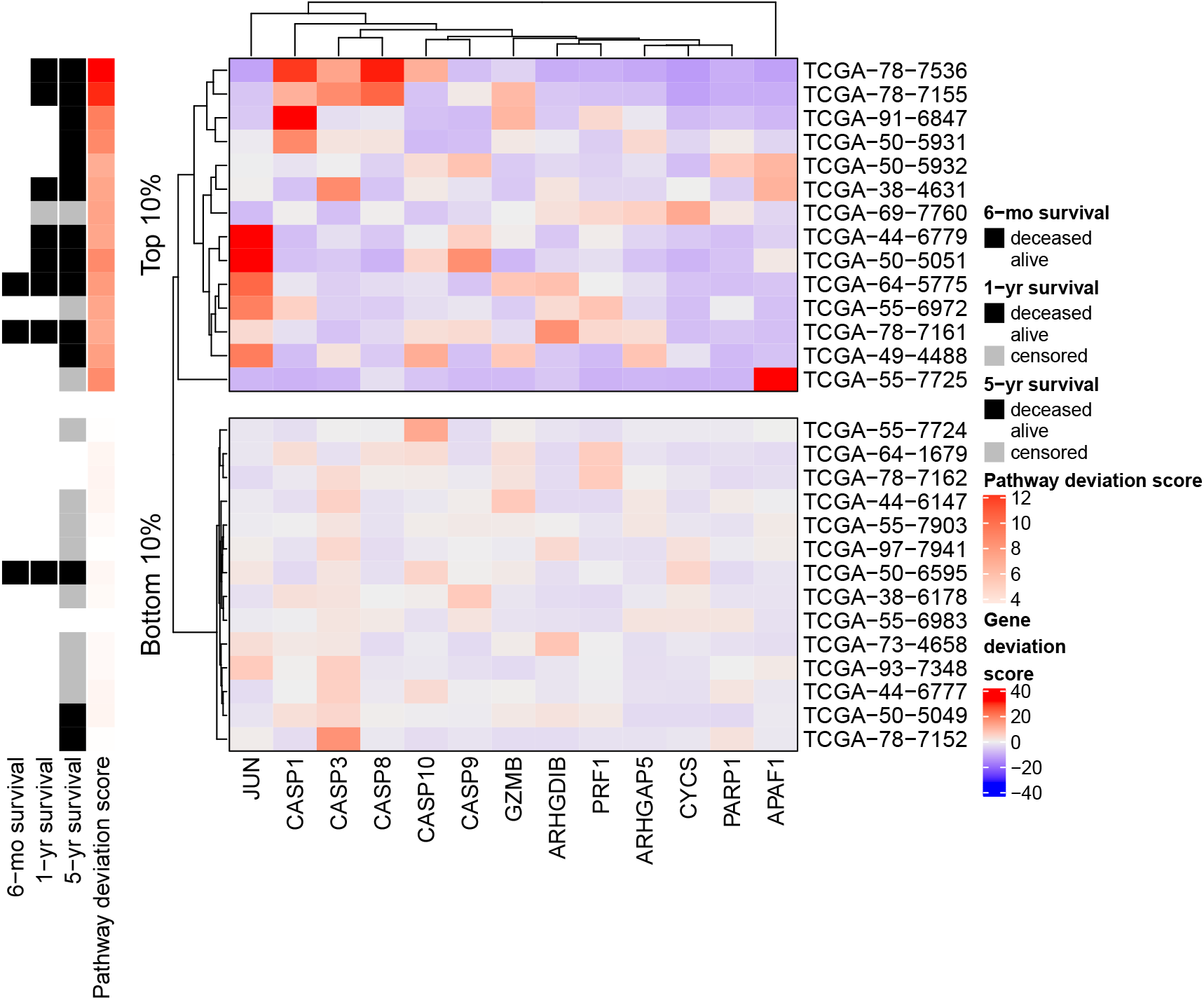
Scaled per-gene deviation scores for the D4-GDI signaling pathway for individuals corresponding to the top and bottom decile of overall pathway deviation scores. Red scores correspond to highly aberrant gene scores with respect to each individual’s global score, while blue indicates gene scores close to the overall population average. Annotations on the left indicate the 6-month, 1-year, and 5-year survival status (deceased, alive, or censored) and overall pathway deviation score for each individual. Genes and individuals within each sub-plot are hierarchically clustered using the Euclidean distance and complete linkage.

To provide an intuitive link between these gene-specific deviation scores with the original batch-corrected multi-omics data that were input into *padma*, we further focus on the three genes (CASP1, CASP3, and CASP8) for which large deviation scores were observed for the two highly aberrant individuals (TCGA-78-7536 and TCGA-78-7155) in the D4-GDI signaling pathway. We plot boxplots of the Z-scores for each available omic for the three genes across all 144 individuals with lung cancer (Figure 3), specifically highlighting the two aforementioned individuals; full plots of all 13 genes in the pathway are included in Supplementary Figure 1. This plot reveals that both individuals are indeed notable for their overexpression, with respect to the other individuals, of miRNA hsa-mir-421 (Figure 3D), which is predicted to target CASP3; consistent with this observation, both individuals had weaker CASP3 expression than average (although we note that its expression was not particularly extreme with respect to the full sample). Individual TCGA-78-7536 appears to have a hypomethylated CASP1 promoter, but a significantly higher number of copies of CASP8, while individual TCGA-78-7155 is characterized by a large underexpression of CASP8 with respect to other individuals. Both individuals appear to have deletions of CASP3, and hypermethylated CASP8 promoters. This seems to indicate that, although the large overall pathway deviations for these two individuals share some common etiologies, each also exhibit unique characteristics.

**Fig. 3.**
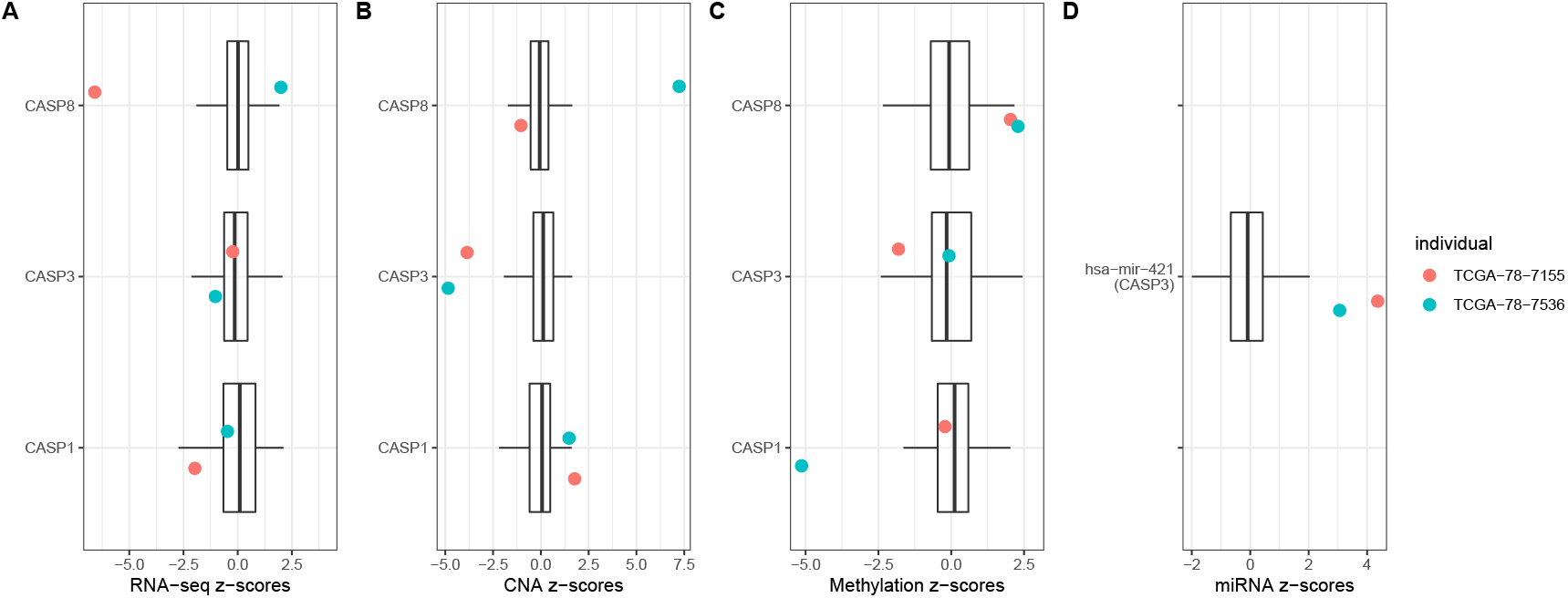
Boxplots of Z-scores of gene expression (A), copy number alterations (B), methylation (C), and miRNA expression (D) for all individuals with lung cancer, with the 3 genes (CASP1, CASP3, CASP8) and one miRNA (hsa-mir-421, predicted to target CASP3) of interest in the D4-GDI signaling pathway. The two individuals with the largest pathway deviation score (TCGA-78-7155, TCGA-78-7536) are highlighted in red and turquoise, respectively.

As overall pathway deviation scores represent the multi-dimensional average of these gene-specific deviation scores, a deeper investigation into them can also provide useful insight for a given pathway. We first note that the distribution of deviation scores for the D4-GDI signaling pathway (Figure 4A) is highly skewed, with a handful of individuals (e.g., TCGA-78-7536, TCGA-78-7155, TCGA-91-6847, TCGA-50-5931, TCGA-50-5051, and TCGA-66-7725) characterized by particularly large scores with respect to the remaining individuals. The individual with the most aberrant score for this pathway, TCGA-78-7536, had a single pathway-specific somatic mutation in the CASP1 gene, and a total of 7 cancer-specific driver gene mutations (corresponding to the 80th percentile of individuals considered here). Although these pathway deviation scores are calculated across all dimensions of the MFA, it can also be useful to represent individuals in the first two components of the consensus MFA space (Figure 4B); the farther away an individual is from the origin over multiple MFA dimensions, the larger the corresponding pathway deviation score. In this case, we see that TCGA-78-7536 is a large positive and negative outlier in the second (9.55% total variance explained), and third (8.07% total variance explained) MFA components, respectively, although less so in the first component (11.97% total variance explained). In addition, we note that RNA-seq is the major driver of the first MFA dimension (54.38% contribution), while promoter methylation and copy number alterations take a larger role in the second and third dimensions (42.29% and 59.18% contribution, respectively). miRNA expression appears to play a fairly minor role in the MFA, with its maximum contribution (21.14%) occurring at only the 16th dimension.

**Fig. 4.**
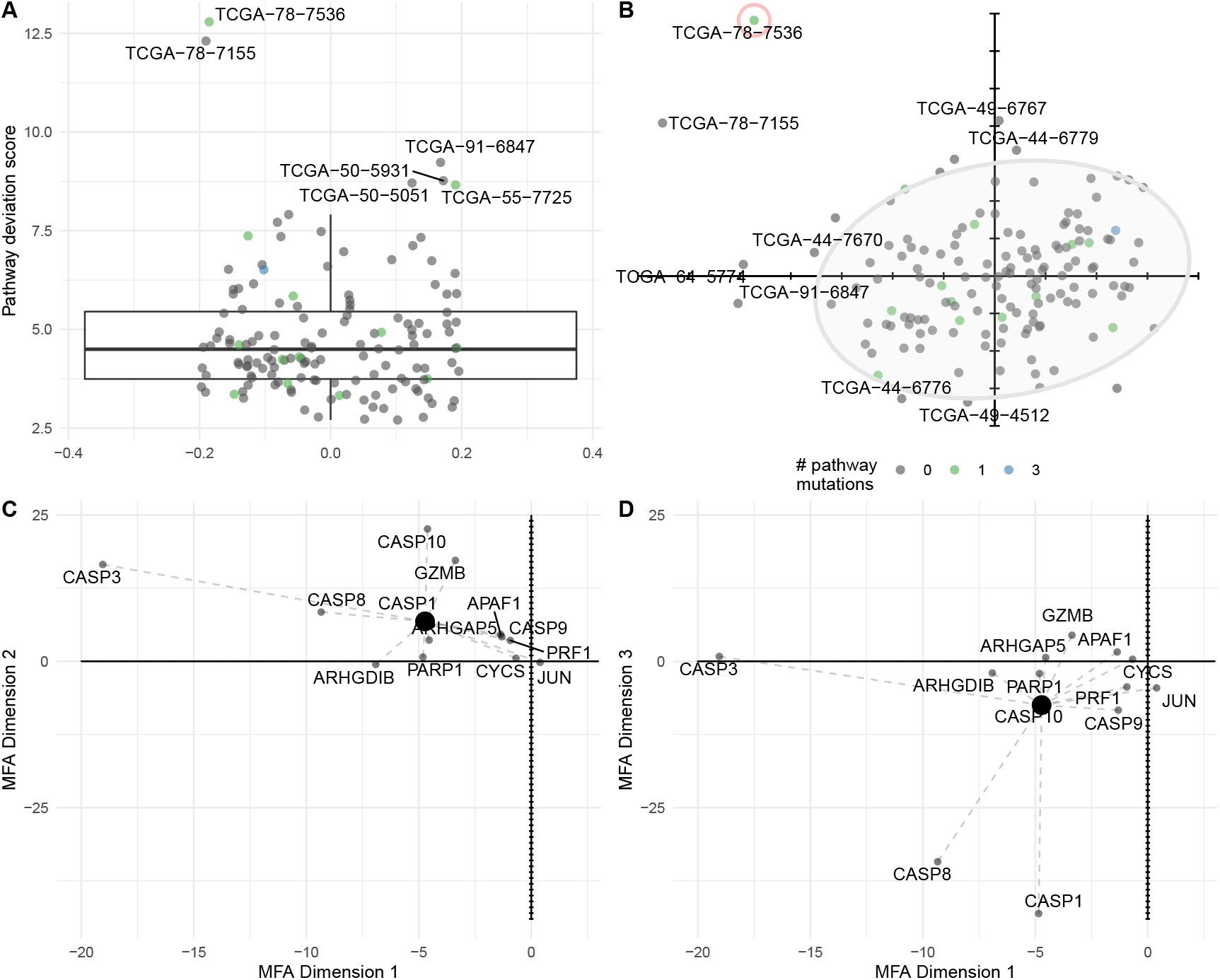
(A) Distribution of pathway deviation scores for the D4-GDI signaling pathway in lung cancer; individuals with unusually large scores are labeled. (B) Factor map, representing the first two components of the MFA for the D4-GDI signaling pathway in lung cancer, with normal confidence ellipse superimposed. Individuals with extreme values in each plot are labeled with their barcode identifiers and colored by the number of pathway-specific nonsynonymous mutations. For the individual circled in red, TCGA-78-7536, a partial factor map representing the first MFA components 1 and 2 is plotted in (C), and MFA components 1 and 3 in (D). The large black dot represents the individual’s overall pathway deviation score, as plotted in panel (B) for the first two axes, and gene-specific scores are joined to this point with dotted lines.

When examining the partial factor maps for this individual over the first three MFA dimensions (Figures 4C-D), we note the large contribution of CASP3 (axis 1), CASP10 (axis 2), CASP1 and CASP 8 (axis 3), as evidenced by their distance from the origin in these dimensions. Overall, this is consistent with the previous gene-level analyses (Figure 2), where hypomethylation in CASP1 and large copy number gains for CASP3 and CASP8 with respect to the population were identified for this individual. Other individuals with large overall deviation scores (e.g., TCGA-50-5931) are not obvious outliers in the first two MFA dimensions, reflecting the fact that additional dimensions play a more important role for them. Taken together, the individualized gene-specific and overall pathway deviation scores output by *padma* provide complementary and interesting exploratory insight into atypical multi-omic profiles for a given pathway of interest (here, the D4-GDI signaling pathway in lung cancer).

### 3.5 Pathway deviation scores globally recapitulate histological grade in breast cancer

For some cancers, additional clinical phenotypes beyond survival information may be of particular interest; to illustrate the use of *padma* in such a case, we focus on histological grade for breast cancer. To quantify whether pathway deviation scores tend to be associated with histological grade in breast cancer, we performed a one-way ANOVA on the three measures that comprise histological grade for each of the 1136 pathways. Based on the BH-adjusted p-values from an F-test (FDR < 5%), all (1136) or nearly all (1135) pathways were found to have deviation scores that are significantly correlated with mitotic index and nuclear pleomorphism. Intriguingly, no pathways were found to be associated with degree of glandular/tubule formation; this may in part be due to the large proportion of individuals identified as grade III (poorly differentiated) for this measure (n = 285). The rankings of pathways based on mitotic index and nuclear pleomorphism were generally in agreement (Supplementary Figure 2). In all but two cases, higher deviation pathway scores corresponded to the higher grades for these two measures, corresponding to more aggressive tumors; the two exceptions were the Presynaptic nicotinic acetylcholine receptor and Highly calcium permeable postsynaptic nicotinic acetylcholine receptor pathways (both from Reactome), for which the largest pathway deviation scores were associated with grade II, rather than grade III, of the mitotic index.

To prioritize pathways among this list, we calculated the rank product of the individual rankings by p-value for mitosis and nuclear pleomorphism; the top 10 pathways according to this joint ranking are shown in Supplementary Table 2 (see Supplementary Table 5 for the full gene lists in each pathway). The Wnt signaling pathway, which is made up of 63 genes, had the highest combined ranking for these two histological measures. Of this set of genes, all had RNA-seq, methylation, and CNA measures available, with the exception of FAM123B and PSMD10 (no CNA measures with nonzero variance) and PSMB1 to PSMB10, PSMC2, PSMC3, PSMC5, PSMC6, PSME1, and PSME2 (no promoter methylation measures). miRNA-seq data were included for only two predicted target pairs: hsa-mir-375 → CTNNB1 and hsa-mir-320a → CTNNB1. Over the 63 genes in the pathway, 453 individuals had no nonsynonymous mutations, while 39, 6, 3, 2, and 1 individuals had 1, 2, 3, 4, or 5 such mutations; APC, PSMD1, and FAM123B were most often characterized by mutations (10, 7, and 7 individuals affected, respectively).

Similarly to the distribution of D4-GDI pathway scores in lung adenocarcinomas, a small number of breast cancer patients are characterized by highly aberrant scores in the Wnt signaling pathway, including TCGA-BH-A1FM, TCGA-E9-A22G, and TCGA-EW-A1PH, and the number of pathway-specific nonsynonymous somatic mutations does not appear to be related to this score. The associated factor map on the first two dimensions of the MFA (Figure 5A) clearly captures relevant biological structure from the data, as evidenced by the quasi-separation of individuals in different intrinsic inferred molecular subtypes (AIMS). Notably, individuals with Basal and Luminal A breast cancer are clearly separated in the first two dimensions and tend to respectively have positive and negative loadings in the first dimension of the MFA; Luminal B and Normal-like subtypes largely overlap with the Luminal A subtype for this pathway, while Her2 is located intermediate to the Luminal and Basal subtypes, as could be anticipated due to the equal prevalence of Her2 amplification in both Luminal and Basal subtypes. Similar relevant biological signal can be seen when considering a larger spectrum of pathways (Figure 5C). In particular, individuals with the Basal and Luminal B subtypes tend to have much more highly variant deviation scores across all pathways, whereas Luminal A and Normal-like subtypes are generally much less variant.

**Fig. 5.**
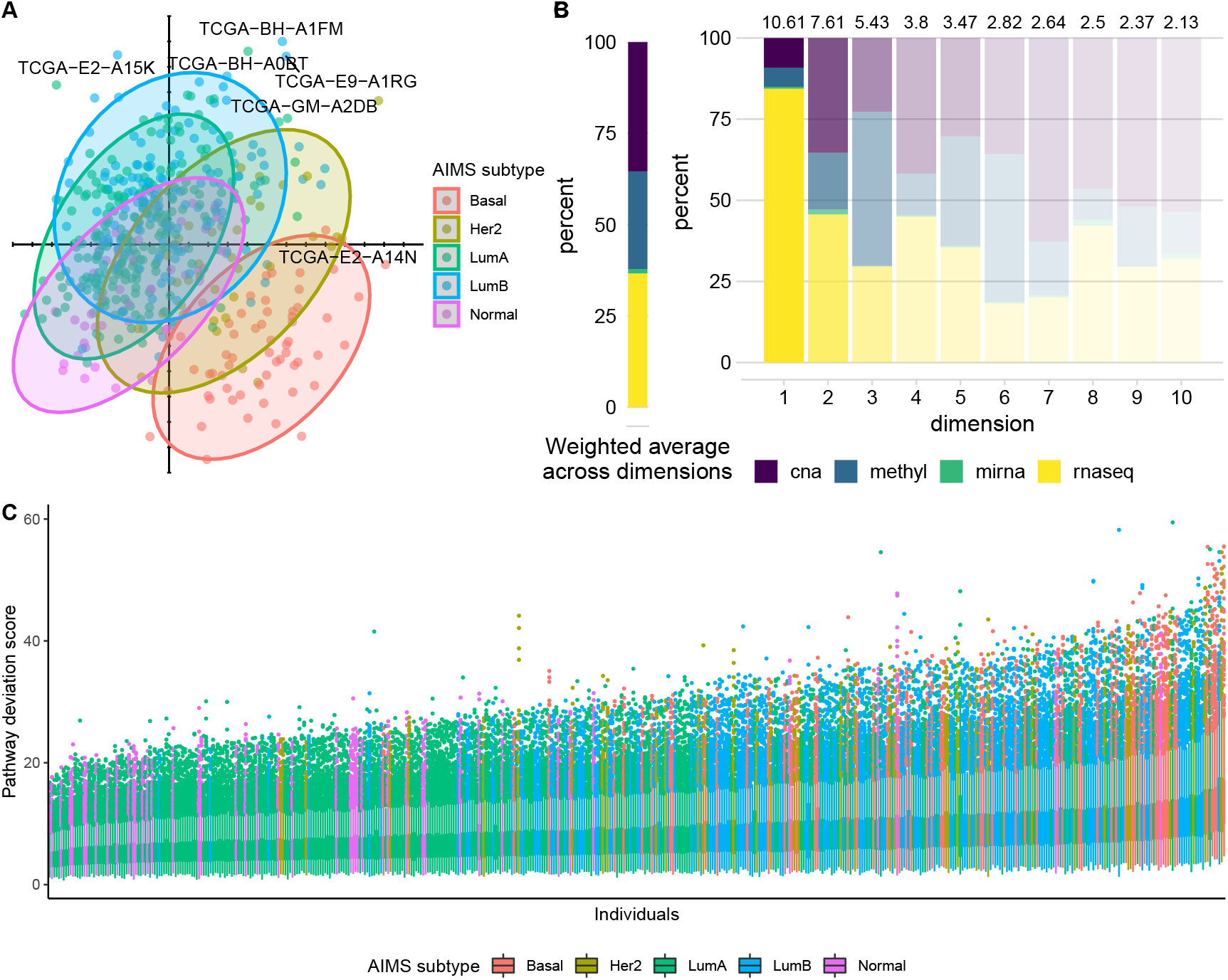
(A) Factor map of individuals, representing the first two components of the MFA, for the Wnt signaling pathway in breast cancer, with normal confidence ellipses superimposed for the five AIMS subtypes. B) Weighted overall percent contribution per omic (left) and for each of the first 10 MFA components (right) for the Wnt signaling pathway, with colors faded according to the percent variance explained for each (represented in text above each bar). (C) Distribution of pathway deviation scores for each individual in the breast cancer data, with individuals colored according to their AIMS subtype.

When examining the percent contribution of each omic to the axes of the MFA for the Wnt signaling pathway (Figure 5B), we remark the preponderant contribution of gene expression to the first component (84.40%), while variability in the second component is largely driven by both gene expression and copy numbers (45.66 and 35.37%, respectively). The large role played by RNA-seq here is consistent with the definition of the AIMS subtypes themselves, which are defined on the basis of gene expression. On average, after weighting by the eigenvalue of each component, gene expression and copy number alterations were found to have similar contributions to the overall variation (36.6%, 35.4%, respectively), while methylation played a less important role (26.8%). For this pathway, as for most others we studied (Supplementary Figure 6), miRNA expression contributed relatively little to the overall variation (1.2%). We do need to be cautious in the interpretation of this phenomenon, as it may be due to real biology or to the mapping uncertainty and much smaller number (with respect to the other omics) of miRNA features (Supplementary Figure 10). Because we have structured the data into gene-tables (and not omics-tables) in *padma*, the MFA weighting leads to a balancing of contributions among genes (and not omics); as such, the drastically smaller number of miRNA features is likely directly linked to the overall smaller contributions to the variance explained.

Taken together, these results illustrate that the *padma* approach, which is used in an unsupervised manner on multi-omic cancer data for a given pathway, is able to recapitulate known sample structure in the form of intrinsic tumor subtypes as well as relevant prognostic factors such as histological grade.

## 4. Conclusions

Unsupervised dimension reduction approaches (such as PCA) have been widely used in genetics and genomics for many years, both to identify sample structure and batch effects (Leek *and others*, 2010) and to visualize overall variation in large data (Gautier *and others*, 2010). Here, we present a generalization of this approach to multi-omic data for investigating biological variation at the pathway-level by aggregating across genes, omic-type, and individuals. Compared to single-omics approaches (for instance, running a PCA on RNA-seq data alone), *padma* accommodates multiple omics-sources which, for some sample sets and pathways, account for more than 50% of the overall variation (Figure 5B). Using MFA to partition variance, we construct a clinically relevant pathway disruption score that correlates with survival outcomes in lung cancer patients, and histological grade in breast cancer patients.

Our MFA-based approach allows investigators to (a) identify overall sources of variation (such as batch effects); (b) prioritize high variance pathways defined by variability across subjects; (c) identify aberrant observations (i.e., individuals) within a given pathway; and (d) identify the genes and omics sources that drive these aberrant observations. As with any analysis of omic data we generally recommend that standard quality control analyses be performed (e.g., genomewide PCA of each omic individually as in Supplementary Figures 3-4, boxplots of normalized read count distributions for each sample) to identify any potential technical outliers before running *padma*. Here, we chose to remove known batch effects prior to the *padma* analysis, but in principle the method could be run on uncorrected data and MFA components correlated with undesired batch effects could be identified and removed prior to computing the pathway deviation score. For large, multi-omic data such as TCGA, *padma* allows investigators to summarize overall variation and assist in generating hypotheses for more targeted analyses and follow-up studies. As a case in point, we identified two lung cancer patients with aberrant multi-omic profiles at three CASP genes. With access to the tumor samples and more fine-grained clinical data, future molecular experiments could help to clarify the role (if any) that these genes play in contributing to lung cancer mortality. Although the multi-omic TCGA data considered here were quite large *(n* = 144 and n = 504 matched samples for lung and breast cancer, respectively), Thioulouse (2011) recently suggested that descriptive methods like PCA and MFA can be used without limitation on the ratio between the number of samples and number of variables; as such we anticipate that *padma* could be useful even for more modestly sized multi-omic datasets.

There are a number of natural extensions and alternative formulations to our MFA-based approach. If comparisons between sets of individuals (e.g., healthy vs. disease) are of interest, the MFA can be based on one set of samples (e.g., healthy, or a “reference set”), and the other set of samples (e.g., diseased, or a “supplementary set”) can be projected onto this original representation. This is accomplished by centering and scaling supplementary individuals to the same scale as the reference individuals, and projecting these rescaled variables into the reference MFA space. In this setting, the interpretation of pathway deviation scores would no longer correspond to the identification of “aberrant” individuals compared to an overall average, but rather individuals that are most different from the reference set (e.g., the most “diseased” as compared to a healthy reference); this strategy would be similar in spirit to the individualized pathway aberrance score (iPAS) approach, which proposed using accumulated (unmatched) normal samples as a reference set (Ahn *and others*, 2014). There is also no reason to limit this approach to pathways, as the analysis could be performed just once, genome-wide (accordingly, inferences would no longer be applicable to specific pathways). Here, we have structured the data with genes representing data tables and omics representing columns within each table. Alternatively, the data could be reweighted by having omics represented as data tables and genes as columns within each, similar to de Tayrac *and others* (2009). Extensions to our work could include incorporating the hierarchical structure of genes within pathways, or relatedness structure among samples. In principle, other types of omics that do not map to genes or pathways (e.g., genotypes on single nucleotide polymorphisms) could also be incorporated. Finally, though we illustrate the use of *padma* for cancer genomics data, we anticipate that it will be broadly useful to other multi-omic applications in human health or agriculture.

## 5. Supplementary Material

Supplementary material, including Supplementary Methods, Tables, and Figures, is available online at http://biostatistics.oxfordjournals.org.

### Funding

This research was supported by the AgreenSkills+ fellowship program, which received funding from the EU’s Seventh Framework Program under grant agreement FP7-60939 (AgreenSkills+ contract).

## Acknowledgments

*Conflict of Interest:* None declared.

## References

Abdi, Hervé, Williams, Lynne J. and Valentin, Domininique. (2013, March). Multiple factor analysis: principal component analysis for multitable and multiblock data sets: Multiple factor analysis. Wiley Interdisciplinary Reviews: Computational Statistics 5(2), 149–179.

Ahn, TaeJin, Lee, Eunjin, Huh, Nam and Park, Taesung. (2014, September). Personalized identification of altered pathways in cancer using accumulated normal tissue data. Bioinformatics 30(17), i422–i429.

Argelaguet, Ricard, Velten, Britta, Arnol, Damien, Dietrich, Sascha, Zenz, Thorsten, Marioni, John C, Buettner, Florian, Huber, Wolfgang and Stegle, Oliver. (2018, June). Multi-Omics Factor Analysis—a framework for unsupervised integration of multi-omics data sets. Molecular Systems Biology 14(6).

Benjamini, Yoav and Hochberg, Yosef. (1995). Controlling the False Discovery Rate: A Practical and Powerful Approach to Multiple Testing. Journal of the Royal Statistical Society. Series B (Methodological) 57(1), 289–300.

Benjamini, Y. and Yekutieli, D. (2001). The control of the false discovery rate in multiple testing under dependency. Annals of Statistics 29(4), 1165–1188.

Bosken, Carol H., Wei, Qingyi, Amos, Christopher I. and Spitz, Margaret R. (2002, July). An analysis of DNA repair as a determinant of survival in patients with non-small-cell lung cancer. Journal of the National Cancer Institute 94(14), 1091–1099.

Bourgon, R., Gentleman, R. and Huber, W. (2010, May). Independent filtering increases detection power for high-throughput experiments. Proceedings of the National Academy of Sciences 107(21), 9546–9551.

Chou, Chih-Hung, Shrestha, Sirjana, Yang, Chi-Dung, Chang, Nai-Wen, Lin, Yu-Ling, Liao, Kuang-Wen, Huang, Wei-Chi, Sun, Ting-Hsuan, Tu, Siang-Jyun, Lee, Wei-Hsiang, Chiew, Men-Yee, Tai, Chun-San, Wei, Ting-Yen, Tsai, Tzi-Ren, Huang, Hsin-Tzu, Wang, Chung-Yu, Wu, Hsin-Yi, Ho, Shu-Yi, Chen, Pin-Rong, Chuang, Cheng-Hsun, Hsieh, Pei-Jung, Wu, Yi-Shin, Chen, Wen-Liang, Li, Meng-Ju, Wu, Yu-Chun, Huang, Xin-Yi, Ng, Fung Ling, Buddhakosai, Waradee, Huang, Pei-Chun, Lan, Kuan-Chun, Huang, Chia-Yen, Weng, Shun-Long, Cheng, Yeong-Nan, Liang, Chao, Hsu, Wen-Lian and others. (2018, January). miRTarBase update 2018: a resource for experimentally validated microRNA-target interactions. Nucleic Acids Research 46(D1), D296–D302.

de Tayrac, Marie, Le, Sebastien, Aubry, Marc, Mosser, Jean and Husson, Francois. (2009). Simultaneous analysis of distinct Omics data sets with integration of biological knowledge: Multiple Factor Analysis approach. BMC Genomics 10(1), 32.

Drier, Y., Sheffer, M. and Domany, E. (2013, April). Pathway-based personalized analysis of cancer. Proceedings of the National Academy of Sciences 110(16), 6388–6393.

Escofier, Brigitte and Pagès, Jérôme. (2014). Analyses factorielles simples et multiples: objectifs, méthodes et interprétation. OCLC: 1040936107.

Gautier, M., Laloë, D.s and Moazami-Goudarzi, K. (2010). Insights into the genetic history of french cattle from dense snp data on 47 worldwide breeds. PloS One 5(9).

Gonzalez-Perez, Abel, Perez-Llamas, Christian, Deu-Pons, Jordi, Tamborero, David, Schroeder, Michael P, Jene-Sanz, Alba, Santos, Alberto and Lopez-Bigas, Nuria. (2013, November). IntOGen-mutations identifies cancer drivers across tumor types. Nature Methods 10(11), 1081–1082.

Gu, Zuguang, Eils, Roland and Schlesner, Matthias. (2016, September). Complex heatmaps reveal patterns and correlations in multidimensional genomic data. Bioinformatics 32(18), 2847–2849.

Heng, Yujing J, Lester, Susan C, Tse, Gary MK, Factor, Rachel E, Allison, Kimberly H, Collins, Laura C, Chen, Yunn-Yi, Jensen, Kristin C, Johnson, Nicole B, Jeong, Jong Cheol, Punjabi, Rahi, Shin, Sandra J, Singh, Kamaljeet, Krings, Gregor, Eberhard, David A, Tan, Puay Hoon, Korski, Konstanty, Waldman, Frederic M, Gutman, David A, Sanders, Melinda, Reis-Filho, Jorge S, Flanagan, Sydney R, Gendoo, Deena MA, Chen, Gregory M, Haibe-Kains, Benjamin, Ciriello, Giovanni, Hoadley, Katherine A, Perou, Charles M and others. (2017, February). The molecular basis of breast cancer pathological phenotypes: Molecular basis of breast cancer pathological phenotypes. The Journal of Pathology 241(3), 375–391.

Husson, François, Lê, Sébastien and Pagès, Jérôme. (2017). Exploratory multivariate analysis by example using R, Second edition edition. Boca Raton: CRC Press.

Josse, J. and Husson, F. (2016). missMDA: a package for handling missing values in multivariate data analysis. Journal of Statistical Software 70(1), 1–31.

Leek, Jeffrey T., Scharpf, Robert B., Bravo, Héctor Corrada, Simcha, David, Langmead, Benjamin, Johnson, W. Evan, Geman, Donald, Baggerly, Keith and Irizarry, Rafael A. (2010). Tackling the widespread and critical impact of batch effects in high-throughput data. Nature Reviews. Genetics 11(10), 733–739.

Liberzon, A., Subramanian, A., Pinchback, R., Thorvaldsdottir, H., Tamayo, P. and Mesirov, J. P. (2011, June). Molecular signatures database (MSigDB) 3.0. Bioinformatics 27(12), 1739–1740.

Liu, Jianfang, Lichtenberg, Tara, Hoadley, Katherine A., Poisson, Laila M., Lazar, Alexander J., Cherniack, Andrew D., Kovatich, Albert J., Benz, Christopher C., Levine, Douglas A., Lee, Adrian V., Omberg, Larsson, Wolf, Denise M., Shriver, Craig D., Thorsson, Vesteinn, Hu, Hai, Caesar-Johnson, Samantha J., Demchok, John A., Felau, Ina, Kasapi, Melpomeni, Ferguson, Martin L., Hutter, Carolyn M., Sofia, Heidi J., Tarnuzzer, Roy, Wang, Zhining, Yang, Liming, Zenklusen, Jean C., Zhang, Jiashan (Julia), Chudamani, Sudha, Liu, Jia, Lolla, Laxmi, Naresh, Rashi, Pihl, Todd, Sun, Qiang, Wan, Yunhu, Wu, Ye, Cho, Juok, DeFreitas, Timothy, Frazer, Scott, Gehlenborg, Nils, Getz, Gad, Heiman, David I., Kim, Jaegil, Lawrence, Michael S., Lin, Pei, Meier, Sam, Noble, Michael S., Saksena, Gordon, Voet, Doug, Zhang, Hailei, Bernard, Brady, Chambwe, Nyasha, Dhankani, Varsha, Knijnenburg, Theo, Kramer, Roger, Leinonen, Kalle, Liu, Yuexin, Miller, Michael, Reynolds, Sheila, Shmulevich, Ilya, Thorsson, Vesteinn, Zhang, Wei, Akbani, Rehan, Broom, Bradley M., Hegde, Apurva M., Ju, Zhenlin, Kanchi, Rupa S., Korkut, Anil, Li, Jun, Liang, Han, Ling, Shiyun, Liu, Wenbin, Lu, Yiling, Mills, Gordon B., Ng, Kwok-Shing, Rao, Arvind, Ryan, Michael, Wang, Jing, Weinstein, John N., Zhang, Jiexin, Abeshouse, Adam, Armenia, Joshua, Chakravarty, Debyani, Chatila, Walid K., de Bruijn, Ino, Gao, Jianjiong, Gross, Benjamin E., Heins, Zachary J., Kundra, Ritika, La, Konnor, Ladanyi, Marc, Luna, Augustin, Nissan, Moriah G., Ochoa, Angelica, Phillips, Sarah M., Reznik, Ed, Sanchez-Vega, Francisco, Sander, Chris, Schultz, Nikolaus, Sheridan, Robert, Sumer, S. Onur, Sun, Yichao, Taylor, Barry S., Wang, Jioajiao, Zhang, Hongxin, Anur, Pavana, Peto, Myron, Spellman, Paul, Benz, Christopher, Stuart, Joshua M., Wong, Christopher K., Yau, Christina, Hayes, D. Neil, Parker, Joel S., Wilkerson, Matthew D., Ally, Adrian, Balasundaram, Miruna, Bowlby, Reanne, Brooks, Denise, Carlsen, Rebecca, Chuah, Eric, Dhalla, Noreen, Holt, Robert, Jones, Steven J.M., Kasaian, Katayoon, Lee, Darlene, Ma, Yussanne, Marra, Marco A., Mayo, Michael, Moore, Richard A., Mungall, Andrew J., Mungall, Karen, Robertson, A. Gordon, Sadeghi, Sara, Schein, Jacqueline E., Sipahimalani, Payal, Tam, Angela, Thiessen, Nina, Tse, Kane, Wong, Tina, Berger, Ashton C., Beroukhim, Rameen, Cherniack, Andrew D., Cibulskis, Carrie, Gabriel, Stacey B., Gao, Galen F., Ha, Gavin, Meyerson, Matthew, Schumacher, Steven E., Shih, Juliann, Kucherlapati, Melanie H., Kucherlapati, Raju S., Baylin, Stephen, Cope, Leslie, Danilova, Ludmila, Bootwalla, Moiz S., Lai, Phillip H., Maglinte, Dennis T., Van Den Berg, David J., Weisenberger, Daniel J., Auman, J. Todd, Balu, Saianand, Bodenheimer, Tom, Fan, Cheng, Hoadley, Katherine A., Hoyle, Alan P., Jefferys, Stuart R., Jones, Corbin D., Meng, Shaowu, Mieczkowski, Piotr A., Mose, Lisle E., Perou, Amy H., Perou, Charles M., Roach, Jeffrey, Shi, Yan, Simons, Janae V., Skelly, Tara, Soloway, Matthew G., Tan, Donghui, Veluvolu, Umadevi, Fan, Huihui, Hinoue, Toshinori, Laird, Peter W., Shen, Hui, Zhou, Wanding, Bellair, Michelle, Chang, Kyle, Covington, Kyle, Creighton, Chad J., Dinh, Huyen, Doddapaneni, HarshaVardhan, Donehower, Lawrence A., Drummond, Jennifer, Gibbs, Richard A., Glenn, Robert, Hale, Walker, Han, Yi, Hu, Jianhong, Korchina, Viktoriya, Lee, Sandra, Lewis, Lora, Li, Wei, Liu, Xiuping, Morgan, Margaret, Morton, Donna, Muzny, Donna, Santibanez, Jireh, Sheth, Margi, Shinbro, Eve, Wang, Linghua, Wang, Min, Wheeler, David A., Xi, Liu, Zhao, Fengmei, Hess, Julian, Appelbaum, Elizabeth L., Bailey, Matthew, Cordes, Matthew G., Ding, Li, Fronick, Catrina C., Fulton, Lucinda A., Fulton, Robert S., Kandoth, Cyriac, Mardis, Elaine R., McLellan, Michael D., Miller, Christopher A., Schmidt, Heather K., Wilson, Richard K., Crain, Daniel, Curley, Erin, Gardner, Johanna, Lau, Kevin, Mallery, David, Morris, Scott, Paulauskis, Joseph, Penny, Robert, Shelton, Candace, Shelton, Troy, Sherman, Mark, Thompson, Eric, Yena, Peggy, Bowen, Jay, Gastier-Foster, Julie M., Gerken, Mark, Leraas, Kristen M., Lichtenberg, Tara M., Ramirez, Nilsa C., Wise, Lisa, Zmuda, Erik, Corcoran, Niall, Costello, Tony, Hovens, Christopher, Carvalho, Andre L., de Carvalho, Ana C., Fregnani, Jose H., Longatto-Filho, Adhemar, Reis, Rui M., Scapulatempo-Neto, Cristovam, Silveira, Henrique C.S., Vidal, Daniel O., Burnette, Andrew, Eschbacher, Jennifer, Hermes, Beth, Noss, Ardene, Singh, Rosy, Anderson, Matthew L., Castro, Patricia D., Ittmann, Michael, Huntsman, David, Kohl, Bernard, Le, Xuan, Thorp, Richard, Andry, Chris, Duffy, Elizabeth R., Lyadov, Vladimir, Paklina, Oxana, Setdikova, Galiya, Shabunin, Alexey, Tavobilov, Mikhail, McPherson, Christopher, Warnick, Ronald, Berkowitz, Ross, Cramer, Daniel, Feltmate, Colleen, Horowitz, Neil, Kibel, Adam, Muto, Michael, Raut, Chandrajit P., Malykh, Andrei, Barnholtz-Sloan, Jill S., Barrett, Wendi, Devine, Karen, Fulop, Jordonna, Ostrom, Quinn T., Shimmel, Kristen, Wolinsky, Yingli, Sloan, Andrew E., De Rose, Agostino, Giuliante, Felice, Goodman, Marc, Karlan, Beth Y., Hagedorn, Curt H., Eckman, John, Harr, Jodi, Myers, Jerome, Tucker, Kelinda, Zach, Leigh Anne, Deyarmin, Brenda, Hu, Hai, Kvecher, Leonid, Larson, Caroline, Mural, Richard J., Somiari, Stella, Vicha, Ales, Zelinka, Tomas, Bennett, Joseph, Iacocca, Mary, Rabeno, Brenda, Swanson, Patricia, Latour, Mathieu, Lacombe, Louis, Têtu, Bernard, Bergeron, Alain, McGraw, Mary, Staugaitis, Susan M., Chabot, John, Hibshoosh, Hanina, Sepulveda, Antonia, Su, Tao, Wang, Timothy, Potapova, Olga, Voronina, Olga, Desjardins, Laurence, Mariani, Odette, Roman-Roman, Sergio, Sastre, Xavier, Stern, Marc-Henri, Cheng, Feixiong, Signoretti, Sabina, Berchuck, Andrew, Bigner, Darell, Lipp, Eric, Marks, Jeffrey, McCall, Shannon, McLendon, Roger, Secord, Angeles, Sharp, Alexis, Behera, Madhusmita, Brat, Daniel J., Chen, Amy, Delman, Keith, Force, Seth, Khuri, Fadlo, Magliocca, Kelly, Maithel, Shishir, Olson, Jeffrey J., Owonikoko, Taofeek, Pickens, Alan, Ramalingam, Suresh, Shin, Dong M., Sica, Gabriel, Van Meir, Erwin G., Zhang, Hongzheng, Eijckenboom, Wil, Gillis, Ad, Korpershoek, Esther, Looijenga, Leendert, Oosterhuis, Wolter, Stoop, Hans, van Kessel, Kim E., Zwarthoff, Ellen C., Calatozzolo, Chiara, Cuppini, Lucia, Cuzzubbo, Stefania, DiMeco, Francesco, Finocchiaro, Gaetano, Mattei, Luca, Perin, Alessandro, Pollo, Bianca, Chen, Chu, Houck, John, Lohavanichbutr, Pawadee, Hartmann, Arndt, Stoehr, Christine, Stoehr, Robert, Taubert, Helge, Wach, Sven, Wullich, Bernd, Kycler, Witold, Murawa, Dawid, Wiznerowicz, Maciej, Chung, Ki, Edenfield, W. Jeffrey, Martin, Julie, Baudin, Eric, Bubley, Glenn, Bueno, Raphael, De Rienzo, Assunta, Richards, William G., Kalkanis, Steven, Mikkelsen, Tom, Noushmehr, Houtan, Scarpace, Lisa, Girard, Nicolas, Aymerich, Marta, Campo, Elias, Giné, Eva, Guillermo, Armando López, Van Bang, Nguyen, Hanh, Phan Thi, Phu, Bui Duc, Tang, Yufang, Colman, Howard, Evason, Kimberley, Dottino, Peter R., Martignetti, John A., Gabra, Hani, Juhl, Hartmut, Akeredolu, Teniola, Stepa, Serghei, Hoon, Dave, Ahn, Keunsoo, Kang, Koo Jeong, Beuschlein, Felix, Breggia, Anne, Birrer, Michael, Bell, Debra, Borad, Mitesh, Bryce, Alan H., Castle, Erik, Chandan, Vishal, Cheville, John, Copland, John A., Farnell, Michael, Flotte, Thomas, Giama, Nasra, Ho, Thai, Kendrick, Michael, Kocher, Jean-Pierre, Kopp, Karla, Moser, Catherine, Nagorney, David, O’Brien, Daniel, O’Neill, Brian Patrick, Patel, Tushar, Petersen, Gloria, Que, Florencia, Rivera, Michael, Roberts, Lewis, Smallridge, Robert, Smyrk, Thomas, Stanton, Melissa, Thompson, R. Houston, Torbenson, Michael, Yang, Ju Dong, Zhang, Lizhi, Brimo, Fadi, Ajani, Jaffer A., Angulo Gonzalez, Ana Maria, Behrens, Carmen, Bondaruk, Jolanta, Broaddus, Russell, Czerniak, Bogdan, Esmaeli, Bita, Fujimoto, Junya, Gershenwald, Jeffrey, Guo, Charles, Lazar, Alexander J., Logothetis, Christopher, Meric-Bernstam, Funda, Moran, Cesar, Ramondetta, Lois, Rice, David, Sood, Anil, Tamboli, Pheroze, Thompson, Timothy, Troncoso, Patricia, Tsao, Anne, Wistuba, Ignacio, Carter, Candace, Haydu, Lauren, Hersey, Peter, Jakrot, Valerie, Kakavand, Hojabr, Kefford, Richard, Lee, Kenneth, Long, Georgina, Mann, Graham, Quinn, Michael, Saw, Robyn, Scolyer, Richard, Shannon, Kerwin, Spillane, Andrew, Stretch, Jonathan, Synott, Maria, Thompson, John, Wilmott, James, Al-Ahmadie, Hikmat, Chan, Timothy A., Ghossein, Ronald, Gopalan, Anuradha, Levine, Douglas A., Reuter, Victor, Singer, Samuel, Singh, Bhuvanesh, Tien, Nguyen Viet, Broudy, Thomas, Mirsaidi, Cyrus, Nair, Praveen, Drwiega, Paul, Miller, Judy, Smith, Jennifer, Zaren, Howard, Park, Joong-Won, Hung, Nguyen Phi, Kebebew, Electron, Linehan, W. Marston, Metwalli, Adam R., Pacak, Karel, Pinto, Peter A., Schiffman, Mark, Schmidt, Laura S., Vocke, Cathy D., Wentzensen, Nicolas, Worrell, Robert, Yang, Hannah, Moncrieff, Marc, Goparaju, Chandra, Melamed, Jonathan, Pass, Harvey, Botnariuc, Natalia, Caraman, Irina, Cernat, Mircea, Chemencedji, Inga, Clipca, Adrian, Doruc, Serghei, Gorincioi, Ghenadie, Mura, Sergiu, Pirtac, Maria, Stancul, Irina, Tcaciuc, Diana, Albert, Monique, Alexopoulou, Iakovina, Arnaout, Angel, Bartlett, John, Engel, Jay, Gilbert, Sebastien, Parfitt, Jeremy, Sekhon, Harman, Thomas, George, Rassl, Doris M., Rintoul, Robert C., Bifulco, Carlo, Tamakawa, Raina, Urba, Walter, Hayward, Nicholas, Timmers, Henri, Antenucci, Anna, Facciolo, Francesco, Grazi, Gianluca, Marino, Mirella, Merola, Roberta, de Krijger, Ronald, Gimenez-Roqueplo, Anne-Paule, Piché, Alain, Chevalier, Simone, McKercher, Ginette, Birsoy, Kivanc, Barnett, Gene, Brewer, Cathy, Farver, Carol, Naska, Theresa, Pennell, Nathan A., Raymond, Daniel, Schilero, Cathy, Smolenski, Kathy, Williams, Felicia, Morrison, Carl, Borgia, Jeffrey A., Liptay, Michael J., Pool, Mark, Seder, Christopher W., Junker, Kerstin, Omberg, Larsson, Dinkin, Mikhail, Manikhas, George, Alvaro, Domenico, Bragazzi, Maria Consiglia, Cardinale, Vincenzo, Carpino, Guido, Gaudio, Eugenio, Chesla, David, Cottingham, Sandra, Dubina, Michael, Moiseenko, Fedor, Dhanasekaran, Renumathy, Becker, Karl-Friedrich, Janssen, Klaus-Peter, Slotta-Huspenina, Julia, Abdel-Rahman, Mohamed H., Aziz, Dina, Bell, Sue, Cebulla, Colleen M., Davis, Amy, Duell, Rebecca, Elder, J. Bradley, Hilty, Joe, Kumar, Bahavna, Lang, James, Lehman, Norman L., Mandt, Randy, Nguyen, Phuong, Pilarski, Robert, Rai, Karan, Schoenfield, Lynn, Senecal, Kelly, Wakely, Paul, Hansen, Paul, Lechan, Ronald, Powers, James, Tischler, Arthur, Grizzle, William E., Sexton, Katherine C., Kastl, Alison, Henderson, Joel, Porten, Sima, Waldmann, Jens, Fassnacht, Martin, Asa, Sylvia L., Schadendorf, Dirk, Couce, Marta, Graefen, Markus, Huland, Hartwig, Sauter, Guido, Schlomm, Thorsten, Simon, Ronald, Tennstedt, Pierre, Olabode, Oluwole, Nelson, Mark, Bathe, Oliver, Carroll, Peter R., Chan, June M., Disaia, Philip, Glenn, Pat, Kelley, Robin K., Landen, Charles N., Phillips, Joanna, Prados, Michael, Simko, Jeffry, Smith-McCune, Karen, VandenBerg, Scott, Roggin, Kevin, Fehrenbach, Ashley, Kendler, Ady, Sifri, Suzanne, Steele, Ruth, Jimeno, Antonio, Carey, Francis, Forgie, Ian, Mannelli, Massimo, Carney, Michael, Hernandez, Brenda, Campos, Benito, Herold-Mende, Christel, Jungk, Christin, Unterberg, Andreas, von Deimling, Andreas, Bossler, Aaron, Galbraith, Joseph, Jacobus, Laura, Knudson, Michael, Knutson, Tina, Ma, Deqin, Milhem, Mohammed, Sigmund, Rita, Godwin, Andrew K., Madan, Rashna, Rosenthal, Howard G., Adebamowo, Clement, Adebamowo, Sally N., Boussioutas, Alex, Beer, David, Giordano, Thomas, Mes-Masson, Anne-Marie, Saad, Fred, Bocklage, Therese, Landrum, Lisa, Mannel, Robert, Moore, Kathleen, Moxley, Katherine, Postier, Russel, Walker, Joan, Zuna, Rosemary, Feldman, Michael, Valdivieso, Federico, Dhir, Rajiv, Luketich, James, Mora Pinero, Edna M., Quintero-Aguilo, Mario, Carlotti, Carlos Gilberto Jr, Dos Santos, Jose Sebastião, Kemp, Rafael, Sankarankuty, Ajith, Tirapelli, Daniela, Catto, James, Agnew, Kathy, Swisher, Elizabeth, Creaney, Jenette, Robinson, Bruce, Shelley, Carl Simon, Godwin, Eryn M., Kendall, Sara, Shipman, Cassaundra, Bradford, Carol, Carey, Thomas, Haddad, Andrea, Moyer, Jeffey, Peterson, Lisa, Prince, Mark, Rozek, Laura, Wolf, Gregory, Bowman, Rayleen, Fong, Kwun M., Yang, Ian, Korst, Robert, Rathmell, W. Kimryn, Fantacone-Campbell, J. Leigh, Hooke, Jeffrey A., Kovatich, Albert J., Shriver, Craig D., DiPersio, John, Drake, Bettina, Govindan, Ramaswamy, Heath, Sharon, Ley, Timothy, Van Tine, Brian, Westervelt, Peter, Rubin, Mark A., Lee, Jung Il, Aredes, Natália D. and others. (2018, April). An Integrated TCGA Pan-Cancer Clinical Data Resource to Drive High-Quality Survival Outcome Analytics. Cell 173(2), 400–416.e11.

Kalkanis, Steven, Mikkelsen, Tom, Noushmehr, Houtan, Scarpace, Lisa, Girard, Nicolas, Aymerich, Marta, Campo, Elias, Gine, Eva, Guillermo, Armando Lopez, Van Bang, Nguyen, Hanh, Phan Thi, Phu, Bui Duc, Tang, Yufang, Colman, Howard, Evason, Kimberley, Dottino, Peter R., Mar-tignetti, John A., Gabra, Hani, Juhl, Hartmut, Akeredolu, Teniola, Stepa, Serghei, Hoon, Dave, Ahn, Keunsoo, Kang, Koo Jeong, Beuschlein, Felix, Breg-gia, Anne, Birrer, Michael, Bell, Debra, Borad, Mitesh, Bryce, Alan H., Castle, Erik, Chandan, Vishal, Cheville, John, Copland, John A., Farnell, Michael, Flotte, Thomas, Giama, Nasra, Ho, Thai, Kendrick, Michael, Kocher, Jean-Pierre, Kopp, Karla, Moser, Catherine, Nagorney, David, O’Brien, Daniel, O’Neill, Brian Patrick, Patel, Tushar, Petersen, Gloria, Que, Florencia, Rivera, Michael, Roberts, Lewis, Smallridge, Robert, Smyrk, Thomas, Stanton, Melissa, Thompson, R. Houston, Torbenson, Michael, Yang, Ju Dong, Zhang, Lizhi, Brimo, Fadi, Ajani, Jaffer A., Angulo Gonzalez, Ana Maria, Behrens, Carmen, Bondaruk, Jolanta, Broaddus, Russell, Czerniak, Bogdan, Esmaeli, Bita, Fujimoto, Junya, Gershenwald, Jeffrey, Guo, Charles, Lazar, Alexander J., Logothetis, Christopher, Meric-Bernstam, Funda, Moran, Cesar, Ra-mondetta, Lois, Rice, David, Sood, Anil, Tamboli, Pheroze, Thompson, Timothy, Troncoso, Patricia, Tsao, Anne, Wistuba, Ignacio, Carter, Candace, Haydu, Lauren, Hersey, Peter, Jakrot, Valerie, Kakavand, Hojabr, Kefford, Richard, Lee, Kenneth, Long, Georgina, Mann, Graham, Quinn, Michael, Saw, Robyn, Scolyer, Richard, Shannon, Kerwin, Spillane, Andrew, Stretch, Jonathan, Syn-ott, Maria, Thompson, John, Wilmott, James, Al-Ahmadie, Hikmat, Chan, Timothy A., Ghossein, Ronald, Gopalan, Anuradha, Levine, Douglas A., Reuter, Victor, Singer, Samuel, Singh, Bhuvanesh, Tien, Nguyen Viet, Broudy, Thomas, Mirsaidi, Cyrus, Nair, Praveen, Drwiega, Paul, Miller, Judy, Smith, Jennifer, Zaren, Howard, Park, Joong-Won, Hung, Nguyen Phi, Kebebew, Electron, Linehan, W. Marston, Metwalli, Adam R., Pacak, Karel, Pinto, Peter A., Schiff-man, Mark, Schmidt, Laura S., Vocke, Cathy D., Wentzensen, Nicolas, Worrell, Robert, Yang, Hannah, Moncrieff, Marc, Goparaju, Chandra, Melamed, Jonathan, Pass, Harvey, Botnariuc, Natalia, Caraman, Irina, Cernat, Mircea, Chemencedji, Inga, Clipca, Adrian, Doruc, Serghei, Gorincioi, Ghenadie, Mura, Sergiu, Pirtac, Maria, Stancul, Irina, Tcaciuc, Diana, Albert, Monique, Alex-opoulou, Iakovina, Arnaout, Angel, Bartlett, John, Engel, Jay, Gilbert, Sébastien, Parfitt, Jeremy, Sekhon, Harman, Thomas, George, Rassl, Doris M., Rintoul, Robert C., Bifulco, Carlo, Tamakawa, Raina, Urba, Walter, Hayward, Nicholas, Timmers, Henri, Antenucci, Anna, Facciolo, Francesco, Grazi, Gianluca, Marino, Mirella, Merola, Roberta, de Krijger, Ronald, Gimenez-Roqueplo, Anne-Paule, Piché, Alain, Chevalier, Simone, McKercher, Ginette, Birsoy, Kivanc, Barnett, Gene, Brewer, Cathy, Farver, Carol, Naska, Theresa, Pennell, Nathan A., Raymond, Daniel, Schilero, Cathy, Smolenski, Kathy, Williams, Felicia, Morrison, Carl, Borgia, Jeffrey A., Liptay, Michael J., Pool, Mark, Seder, Christopher W., Junker, Kerstin, Omberg, Larsson, Dinkin, Mikhail, Manikhas, George, Alvaro, Domenico, Bragazzi, Maria Consiglia, Cardinale, Vincenzo, Carpino, Guido, Gaudio, Eugenio, Chesla, David, Cottingham, Sandra, Dubina, Michael, Moiseenko, Fedor, Dhanasekaran, Renumathy, Becker, Karl-Friedrich, Janssen, Klaus-Peter, Slotta-Huspenina, Julia, Abdel-Rahman, Mohamed H., Aziz, Dina, Bell, Sue, Cebulla, Colleen M., Davis, Amy, Duell, Rebecca, Elder, J. Bradley, Hilty, Joe, Kumar, Bahavna, Lang, James, Lehman, Norman L., Mandt, Randy, Nguyen, Phuong, Pilarski, Robert, Rai, Karan, Schoenfield, Lynn, Senecal, Kelly, Wakely, Paul, Hansen, Paul, Lechan, Ronald, Powers, James, Tischler, Arthur, Grizzle, William E., Sexton, Katherine C., Kastl, Alison, Henderson, Joel, Porten, Sima, Waldmann, Jens, Fassnacht, Martin, Asa, Sylvia L., Schadendorf, Dirk, Couce, Marta, Graefen, Markus, Huland, Hartwig, Sauter, Guido, Schlomm, Thorsten, Simon, Ronald, Tennstedt, Pierre, Olabode, Oluwole, Nelson, Mark, Bathe, Oliver, Carroll, Peter R., Chan, June M., Disaia, Philip, Glenn, Pat, Kelley, Robin K., Landen, Charles N., Phillips, Joanna, Prados, Michael, Simko, Jeffry, Smith-McCune, Karen, VandenBerg, Scott, Roggin, Kevin, Fehrenbach, Ashley, Kendler, Ady, Sifri, Suzanne, Steele, Ruth, Jimeno, Antonio, Carey, Francis, Forgie, Ian, Man-nelli, Massimo, Carney, Michael, Hernandez, Brenda, Campos, Benito, Herold-Mende, Christel, Jungk, Christin, Unterberg, Andreas, von Deimling, Andreas, Bossler, Aaron, Galbraith, Joseph, Jacobus, Laura, Knudson, Michael, Knutson, Tina, Ma, Deqin, Milhem, Mohammed, Sigmund, Rita, Godwin, Andrew K., Madan, Rashna, Rosenthal, Howard G., Adebamowo, Clement, Adebamowo, Sally N., Boussioutas, Alex, Beer, David, Giordano, Thomas, Mes-Masson, Anne-Marie, Saad, Fred, Bocklage, Therese, Landrum, Lisa, Mannel, Robert, Moore, Kathleen, Moxley, Katherine, Postier, Russel, Walker, Joan, Zuna, Rosemary, Feldman, Michael, Valdivieso, Federico, Dhir, Rajiv, Luketich, James, Mora Pinero, Edna M., Quintero-Aguilo, Mario, Carlotti, Carlos Gilberto Jr, Dos Santos, Jose Sebastiao, Kemp, Rafael, Sankarankuty, Ajith, Tirapelli, Daniela, Catto, James, Agnew, Kathy, Swisher, Elizabeth, Creaney, Jenette, Robinson, Bruce, Shelley, Carl Simon, Godwin, Eryn M., Kendall, Sara, Shipman, Cassaundra, Bradford, Carol, Carey, Thomas, Haddad, Andrea, Moyer, Jeffey, Peterson, Lisa, Prince, Mark, Rozek, Laura, Wolf, Gregory, Bowman, Rayleen, Fong, Kwun M., Yang, Ian, Korst, Robert, Rathmell, W. Kimryn, Fantacone-Campbell, J. Leigh, Hooke, Jeffrey A., Kovatich, Albert J., Shriver, Craig D., DiPer-sio, John, Drake, Bettina, Govindan, Ramaswamy, Heath, Sharon, Ley, Timothy, Van Tine, Brian, Westervelt, Peter, Rubin, Mark A., Lee, Jung Il, Aredes, NatÉLIA D. and others. (2018, April). An Integrated TCGA Pan-Cancer Clinical Data Resource to Drive High-Quality Survival Outcome Analytics. Cell 173(2), 400–416.e11.

Lê, Sébastien, Josse, Julie and Husson, François. (2008). FactoMineR: An *R* Package for Multivariate Analysis. Journal of Statistical Software 25(1).

Meng, Chen, Basunia, Azfar, Peters, Bjoern, Gholami, Amin Moghaddas, Kuster, Bernhard and Culhane, Aedín C. (2019, August). MOGSA: Integrative Single Sample Gene-set Analysis of Multiple Omics Data. Molecular & Cellular Proteomics 18(8 suppl 1), S153–S168.

Meng, Chen, Zeleznik, Oana A., Thallinger, Gerhard G., Kuster, Bernhard, Gholami, Amin M. and Culhane, Aedín C. (2016, July). Dimension reduction techniques for the integrative analysis of multi-omics data. Briefings in Bioinformatics 17(4), 628–641.

Odom, Gabriel J., Ban, Yuguang, Liu, Lizhong, Sun, Xiaodian, Pico, Alexander R., Zhang, Bing, Wang, Lily and Chen, Xi. (2019, April). pathwayPCA: an R package for integrative pathway analysis with modern PCA methodology and gene selection. preprint, Bioinformatics.

Pagès, Jerome. (2015). Multiple factor analysis by example using R, Chapman & Hall/CRC the R series. Boca Raton: CRC Press, Taylor & Francis Group. OCLC: ocn903630995.

Paquet, Eric R. and Hallett, Michael T. (2015, January). Absolute assignment of breast cancer intrinsic molecular subtype. Journal of the National Cancer Institute 107(1), 357.

Rau, Andrea, Flister, Michael, Rui, Hallgeir and Auer, Paul L. (2018, July). Exploring drivers of gene expression in the Cancer Genome Atlas. Bioinformatics.

Riffo-Campos, Angela, Riquelme, Ismael and Brebi-Mieville, Priscilla. (2016, December). Tools for Sequence-Based miRNA Target Prediction: What to Choose? International Journal of Molecular Sciences 17(12), 1987.

Singhal, Sunil, Vachani, Anil, Antin-Ozerkis, Danielle, Kaiser, Larry R. and Albelda, Steven M. (2005, June). Prognostic implications of cell cycle, apoptosis, and angiogenesis biomarkers in non-small cell lung cancer: a review. Clinical Cancer Research: An Official Journal of the American Association for Cancer Research 11(11), 3974–3986.

The Cancer Genome Atlas Research Network, Weinstein, John N, Collisson, Eric A, Mills, Gordon B, Shaw, Kenna R Mills, Ozenberger, Brad A, Ellrott, Kyle, Shmulevich, Ilya, Sander, Chris and Stuart, Joshua M. (2013, October). The Cancer Genome Atlas Pan-Cancer analysis project. Nature Genetics 45(10), 1113–1120.

Thioulouse, J. (2011). Simultaneous analysis of a sequence of paired ecological tables: A comparison of several methods. Annals of Applied Statistics 5(4), 2300–2325.

Vaske, Charles J., Benz, Stephen C., Sanborn, J. Zachary, Earl, Dent, Szeto, Christopher, Zhu, Jingchun, Haussler, David and Stuart, Joshua M. (2010, June). Inference of patient-specific pathway activities from multi-dimensional cancer genomics data using PARADIGM. Bioinformatics 26(12), i237–i245.

Verbeke, Lieven P. C., Van den Eynden, Jimmy, Fierro, Ana Carolina, Demeester, Piet, Fostier, Jan and Marchal, Kathleen. (2015, July). Pathway Relevance Ranking for Tumor Samples through Network-Based Data Integration. PLOS ONE 10(7), e0133503.

Wold, S., Geladi, P., Esbensen, K. and Ohman, J. (1987). Multi-way principal componentsand PLS-analysis. Journal of Chemometrics 1(1), 41–56.

Zhang, Yaqin, Rivera Rosado, Leslie A., Moon, Sun Young and Zhang, Baolin. (2009, May). Silencing of D4-GDI inhibits growth and invasive behavior in MDA-MB-231 cells by activation of Rac-dependent p38 and JNK signaling. The Journal of Biological Chemistry 284(19), 12956–12965.

Zhang, Yaqin and Zhang, Baolin. (2006, June). D4-GDI, a Rho GTPase regulator, promotes breast cancer cell invasiveness. Cancer Research 66(11), 5592–5598.

