## Supplementary methods + figures + tables for "Individualized multi-omic pathway deviation scores using multiple factor analysis"

**Introduction**

This presentation of the Multiple Factorial Analysis (MFA) is mainly borrowed from Abdi et al. (2013) and Abdi and Williams (2010). MFA applies to the case when several ($K$) sets of variables describe a same set of $I$ individuals (observations). MFA is part of the family of methods based on principal components analysis (PCA) and makes use of the basic notions of inertia and distance.

*Inertia and distance*

To start, consider a single data table $\mathbf{X} = \{x_{ij}\}_{I,J}$. The inertia of $x_j$, the j$^{th}$ column of $\mathbf{X}$, is defined as the sum of its squared elements:

$$\gamma_j^2 = \sum_{i=1}^{I} x_{i,j}^2.$$

When the column is centered, $\gamma_j^2$ corresponds to the variance of $x_j$. The sum of all $\gamma_j^2$ across columns is denoted $\mathcal{I}$ and is called the *inertia* of the data table, or the total inertia.

The center of gravity of the rows, denoted **g**, is the vector of the means of each column of **X**. When **X** is column-centered (which is commonly the case), **g** is equal to the $J$-dimensional null vector. The Euclidean distance of the $i^{th}$ observation to **g** is equal to

$$d_i^2 = \sum_{j=1}^{J} x_{i,j}^2.$$

The sum of all $d_i^2$ is equal to the total inertia $\mathcal{I}$ of the data table.

**Principal components analysis**

PCA computes new variables called *principal components*, which are obtained as linear combinations of the original variables, and is based on the concept of inertia. By design, the first principal component is required to have the largest possible inertia. The second component is then constructed to have the largest possible inertia, under the constraint of orthogonality with respect to the first component; the other components are computed in a similar manner. These new variables are called *factor scores*, which can be interpreted geometrically as the projections of the observations onto the principal components. The coefficients of the linear combinations used to compute the factor scores are called the *loadings*.

Specifically, **X** is factorized via a Singular Value Decomposition (SVD) as $\mathbf{X} = \mathbf{P} \Delta \mathbf{Q}^T$, where **P** is an $I \times L$ matrix of normalized left singular vectors (with $L$ being the rank of **X** ), **Q** the $J \times L$ matrix of normalized right singular vectors, and $\Delta$ the $L \times L$ diagonal matrix of the $L$ ordered singular values ($\beta_{[1]}, \ldots, \beta_{[l]}, .., \beta_{[L]}$), such that $\beta_{[1]} \geq \ldots \geq \beta_{[l]} \geq \ldots \geq \beta_{[L]}$. For convenience, we will denote the largest singular value of **X**, $\beta_{[1]}$, as $\lambda$. Matrices **P** and **Q** are orthonormal matrices (i.e., $\mathbf{Q}^T\mathbf{Q} = \mathbf{P}^T\mathbf{P} = \mathbf{I}$). The SVD is closely related to the canonical decomposition of $\mathbf{XX}^T$, where **P** is the matrix of normalized eigenvectors of $\mathbf{XX}^T$, **Q** is the matrix of normalized eigenvectors of $\mathbf{X}^T\mathbf{X}$, and the singular values are the square root of the eigenvalues of $\mathbf{XX}^T$.

*Loadings and factor scores*

As **Q** is the matrix of loadings, the matrix of factor scores **F** is equal to:

$$\mathbf{F} = \mathbf{P}\Delta = \mathbf{XQ}.$$

*Deviation scores*

We define the *deviation score* of an observation as the distance of its factor scores to the center of the cloud of points; that is, if we note $f_{i,l}$ the factor score of the $i^{th}$ individual for the $l^{th}$ component, the deviation score may be calculated as follows:

$$d_i^2 = \sum_{l=1}^{L} f_{i,l}^2.$$

Note that this is equivalent to the Euclidean distance defined above, assuming **X** has been column-centered.

#### Multiple factor analysis

We now consider a set of $K$ tables describing the same set of $I$ individuals (observations). We denote the merged dataset $\mathbf{X} = (\mathbf{X}_{[1]}, \ldots, \mathbf{X}_{[k]}, \ldots \mathbf{X}_{[K]})$, where $\mathbf{X}_{[k]}$ is the data matrix corresponding to the $k^{th}$ set of variables, possibly preprocessed (generally centered and normalized). MFA comprises two main steps:

1. PCA of each table $\mathbf{X}_{[k]}$.
   In this first step, each table $\mathbf{X}_{[k]}$ is analyzed via a standard PCA. The largest singular value obtained from the SVD of $\mathbf{X}_{[k]}$ is noted $\lambda_{[k]}$. We note **A** the $K \times K$ diagonal matrix whose general term $\mathbf{A}[i,i]$ is equal to $\lambda_{[i]}$.

   The MFA consists of a PCA of the merged table **X**, weighted so that the influence of each table $\mathbf{X}_{[k]}$ is balanced.

2. PCA of the weighted table.
   We note $\mathbf{X}^*$ the set of the merged reweighted $\mathbf{X}^*_{[k]}$ matrices, obtained by dividing all the elements of $\mathbf{X}_{[k]}$ by their largest singular value $\lambda_{[k]}$:

$$\mathbf{X}^* = \mathbf{XA}^{-1} = \left[\frac{\mathbf{X}_{[1]}}{\lambda_{[1]}}, \ldots, \frac{\mathbf{X}_{[k]}}{\lambda_{[k]}}, \ldots \frac{\mathbf{X}_{[K]}}{\lambda_{[K]}}\right].$$

The MFA is then computed as the simple PCA of **X***, where the SVD of **X*** is given by **X*** = **P*Δ*Q***$^{\mathbf{T}}$, with **P***$^{\mathbf{T}}$**P*** = **Q***$^{\mathbf{T}}$**Q*** = **I**.

*Loadings*

Because the matrix **X*** concatenates *K* tables, with each of them respectively comprising $J_{[k]}$ variables, the matrix **Q*** of the left singular vectors can be partitioned in the same way as **X*.** Specifically, **Q*** can be expressed as a column block matrix:

$$\mathbf{Q}^* = \begin{bmatrix} \mathbf{Q}^*_{[1]} \\ \vdots \\ \mathbf{Q}^*_{[k]} \\ \vdots \\ \mathbf{Q}^*_{[K]} \end{bmatrix} = \left[\mathbf{Q}^{*\mathrm{T}}_{[1]}, \dots, \mathbf{Q}^{*\mathrm{T}}_{[k]}, \dots \mathbf{Q}^{*\mathrm{T}}_{[K]}\right]^{\mathrm{T}}.$$

The MFA loadings of the $k^{\text{th}}$ table are then calculated as follows:

$$\mathbf{Q}_{[k]} = \lambda_{[k]} \mathbf{Q}^*_{[k]}.$$

*Factor scores*

The factor scores of the observations represent a compromise (i.e. a common representation) for the set of the *K* matrices. They are obtained by:

$$\mathbf{F} = \mathbf{P}^*\boldsymbol{\Delta}^* = \mathbf{X}\mathbf{A}^{-1}\,\mathbf{Q}.$$

Interestingly, this last equation can be rewritten as follows:

$$\mathbf{F} = \mathbf{X}\,\mathbf{A}^{-1}\mathbf{Q} = \left[\frac{\mathbf{X}_{[1]}}{\lambda_{[1]}}, \dots, \frac{\mathbf{X}_{[k]}}{\lambda_{[k]}}, \dots \frac{\mathbf{X}_{[K]}}{\lambda_{[K]}}\right] \begin{bmatrix} \mathbf{Q}_{[1]} \\ \vdots \\ \mathbf{Q}_{[k]} \\ \vdots \\ \mathbf{Q}_{[K]} \end{bmatrix} = \sum_{\boldsymbol{k}=\mathbf{1}}^{\boldsymbol{K}} \frac{\mathbf{1}}{\lambda_{[k]}} \mathbf{X}_{[k]} \mathbf{Q}_{[k]}.$$

We define the partial factor scores of the $k^{\text{th}}$ table by $\mathbf{F}_{[k]} = K\ \frac{\mathbf{1}}{\lambda_{[k]}}\ \mathbf{X}_{[k]}\mathbf{Q}_{[k]}$. The matrix of MFA factor scores **F** then corresponds to the average of all *K* partial factor scores:

$$\mathbf{F} = \frac{1}{K} \sum_{\boldsymbol{k}=\mathbf{1}}^{\boldsymbol{K}} \mathbf{F}_{[k]}.$$

*Deviation scores*

As in a simple PCA, we define the *deviation score* of an observation as the distance of its factor scores to the center of the cloud of points; that is, if we denote $f_{i,l}$ the MFA factor score of the $i^{\text{th}}$ individual for the $l^{\text{th}}$ component:

$$d_i^2 = \sum_{l=1}^{L} f_{i,l}^2\,.$$

#### Contribution of each table to the deviation scores

It may also be of interest to quantify the individual contributions of each data table to the overall deviation score; in particular, this can help identify which data tables have the greatest influence on an individual's deviation score. As above, we denote **F** the matrix of MFA factor scores and $\mathbf{F}_{[k]}$ the matrix of partial factor scores corresponding to the $k^{th}$ table.
We first note that the overall deviation score $d_i^2$ can also be written as follows:

$$d_i^2 = \sum_{l=1}^{L} f_{i,l}^2 + \sum_{k=1}^{K}\sum_{l=1}^{L} f_{i,l}\,(f_{[k]i,l} - f_{i,l}).$$

Noting that

$$\sum_{k=1}^{K}\sum_{l=1}^{L} f_{i,l}\,(f_{[k]i,l} - f_{i,l}) = \sum_{k=1}^{K}\sum_{l=1}^{L} f_{i,l}^2 - (f_{[k]i,l}\, f_{i,l}) = 0,$$

the contribution of the $k^{th}$ table to the deviation score for individual $i$ is equal to

$$d_{i,k} = \frac{\sum_{l=1}^{L} f_{i,l}\,(f_{[k]i,l} - f_{i,l})}{\sum_{l=1}^{L} f_{i,l}^2}.$$

By construction, the contributions of the $K$ tables to the overall deviation score sum to 0. In addition, per-table contributions can take on both negative and positive values according to the extent to which the table influences the deviation of the overall score from the origin (i.e., the global center of gravity across individuals); large positive values correspond to tables with a large influence on the overall deviation of an individual, while large negative values correspond to tables which tend to be most similar to the global average.

#### Contribution of arbitrary groupings of variables to the deviation scores

It is possible to directly quantify the contribution of individual omics (or in fact any arbitrary grouping of variables or any single variable) to the deviation score; in fact, a strategy similar to that for gene-levels contributions to the overall score can be used. Specifically, let the concatenated set of weighted standardized gene tables **X*** be partitioned into $R$ groups, such that

$$\mathbf{X}^* = \left[\mathbf{X}_{[1]}^*, \dots, \mathbf{X}_{[r]}^*, \dots, \mathbf{X}_{[R]}^*\right].$$

The matrix **Q** can then be partitioned the same way, leading to the following:

$$\mathbf{F} = \mathbf{X}^*\mathbf{Q} = \left[\mathbf{X}^*_{[1]}, \ldots, \mathbf{X}^*_{[r]}, \ldots, \mathbf{X}^*_{[R]}\right] \begin{bmatrix} \mathbf{Q}_{[1]} \\ \vdots \\ \mathbf{Q}_{[r]} \\ \vdots \\ \mathbf{Q}_{[R]} \end{bmatrix} = \sum_{r=1}^{R} \mathbf{X}^*_{[r]}\, \mathbf{Q}_{[r]}.$$

For notational simplicity above, the $R$ groups are noted as being made up of contiguous variables in **X***, but any arbitrary partition of variables (e.g., all measurements for each omic) can be used in a similar way by appropriately permuting the columns and rows of $\mathbf{X}^*$ and $\mathbf{Q}$, respectively, to group the variables as needed. The matrix of MFA factor scores **F** can then be written as

$$\mathbf{F} = \frac{1}{R} \sum_{r=1}^{R} \mathbf{F}_{[r]},$$

where $\mathbf{F}_{[r]}$ denotes the matrix of partial factor scores of partition $r$ such that $\mathbf{F}_{[r]} = R\mathbf{X}^*_{[r]}\, \mathbf{Q}_{[r]}$. As the overall pathway deviation score for individual $i$ can be rewritten as

$$d_i^2 = \sum_{l=1}^{L} f_{i,l}^2 + \sum_{r=1}^{R} \sum_{l=1}^{L} f_{i,l}\, (f_{[r]i,l} - f_{i,l}),$$

the individual contribution of the $r^{\text{th}}$ variable partition to the overall pathway deviation score for individual $i$ is equal to

$$d_{i,r}^2 = \frac{\sum_{l=1}^{L} f_{i,l}\, (f_{[r]i,l} - f_{i,l})}{\sum_{l=1}^{L} f_{i,l}^2}.$$

**Supplementary observations and variables**

The results of the MFA can also be used to compute factor scores, loadings, and distances for new observations (rows) and variables (columns) that were not included in the original analysis. For simplicity, we focus on the use of the former in the following discussion, but analogous calculations can be performed for the latter. In particular, assuming that the supplementary rows are scaled in a manner comparable to the original rows of **X**, we can compute the factor scores $\mathbf{f}_{\text{sup}}$ for a supplementary row, which is represented by the $J$-dimensional vector $\mathbf{r}_{sup}^{T}$. The supplementary factor scores are then computed as $\mathbf{f}_{sup} = \mathbf{r}_{sup}^{T}\mathbf{A}^{-1}\mathbf{Q}$, while the partial factor scores are obtained by :

$$\mathbf{f}_{\text{sup}[k]} = K\, \frac{1}{\lambda_{[k]}} \mathbf{r}_{\text{sup}[k]}^{T}\mathbf{Q}.$$

*Deviation scores*

Similar to the case of observations contained in the original data table, the *deviation score* of a supplementary observation is the distance of its factor scores to the center of the cloud of points. That is, if $fsup_{i,l}$ represents the supplementary factor score of the $i^{\text{th}}$ individual for the $l^{\text{th}}$ component:

$$d_i^2 = \sum_{l=1}^{L} fsup_{i,l}^2 .$$

In addition, the contribution of each data table to this overall deviation score for a supplementary individual can be calculated in a manner analogous to that described above.

#### TCGA data acquisition and pre-processing

Briefly, using *TCGA2STAT* (Wan et al., 2016), we downloaded processed TCGA Level 3 data from the Broad Institute Genome Data Analysis Center (GDAC) Firehose on March 18, 2017 for individuals of self-reported European ancestry for whom gene expression, methylation, copy number alterations (CNA), microRNA (miRNA) abundance, and somatic mutation data were all available; this ancestry filter was applied to minimize population-specific variance and focus on the group with the largest available sample size. In addition, two individuals from the BRCA dataset (TCGA-E9-A245, TCGA-BH-A1ES) were identified as outliers with consistently extreme deviation scores across multiple pathways and were removed from the remainder of the analyses; the final sample sizes were thus $n$=504 and $n$=144 individuals for the BRCA and LUAD datasets, respectively.

Per-gene normalized expression estimates were calculated using RSEM (Li et al., 2011). Methylation was quantified using the maximally variant probe from the Illumina Infinium Human Methylation450 BeadChip located within ± 1500bp of the transcription start site, and representative probe beta measures were transformed to the logit scale. Somatic CNAs were called by comparing Affymetrix 6.0 probe intensities from normal (i.e., non-cancer tissue) and cancer tissue, and genome segments were aggregated to gene-level measures by *TCGA2STAT* and *CNTools*. Individuals were classified as carriers or noncarriers of a nonsynonymous somatic mutation for each gene using *TCGA2STAT*. Normalized miRNA abundance was quantified as Reads per million microRNA mapped (RPMMM) values. RNA-seq and miRNA-seq quantifications were TMM-normalized (Robinson and Oshlack, 2010), converted to counts per million (CPM), and $\log_2$-transformed. Only genes with available RNA-seq expression measures were retained for the remainder of the analysis, corresponding to 20,501 and 19,971 genes for BRCA and LUAD, respectively. Finally, batch effects have been shown to have a strong impact on the analysis of high-throughput data in general (Leek et al., 2010) and for the TCGA data specifically (Akulenko et al., 2016). As specific sample plates have been shown to represent significant batch effects in previous analyses (https://bioinformatics.mdanderson.org/BatchEffectsViewer/), each processed omic (with the exception of somatic mutation data) was individually batch adjusted for each cancer to correct for plate-specific effects using `removeBatchEffects` in *limma* (Ritchie et al., 2015). Plots of the first two components from a transcriptome-wide and genome-wide single-omics PCA and multi-omics MFA for the batch-corrected data are included in Supplementary Figures 3 and 4.

#### Choice of curated pathway collection

We consider the pathways included in the MSigDB canonical pathways curated gene set catalog (Liberzon et al., 2011) which includes genes whose products are involved in metabolic and signaling pathways reported in curated public databases. We specifically use the "C2 curated gene sets" catalog from MSigDB v5.2 available at http://bioinf.wehi.edu.au/software/MSigDB as described in the *limma* Bioconductor package (Ritchie et al., 2015). We focus in particular on a collection of 1322 gene sets from public databases, including Biocarta, Pathway Interaction Database (Schaefer et al., 2009), Reactome (Fabregat et al., 2018), Sigma Aldrich, Signaling Gateway, Signal Transduction Knowledge Environment, and the Matrisome Project (Naba et al., 2012), the smallest and largest of which were respectively made up of 6 and 478 genes (median size 29 genes). For the subsequent *padma* analysis, we excluded gene sets for which fewer than 3 genes mapped to quantified features in the TCGA gene expression data, corresponding to a total of 1136 gene sets.

**Details of simulation study**

It is of particular interest to investigate the conditions under which *padma* can correctly identify outlier individuals and the corresponding genes driving their aberrant multi-omic profiles, as well as how it compares to other alternatives. To evaluate the performance of *padma* in a variety of scenarios, we conducted a simulation study as follows.

Assuming a pathway of fixed size with $K$ = 29 genes (the median pathway size of the MSigDB collection) and multi-omic data made up of three assays, we simulated 29 three-dimensional tables for $n$ individuals, $\mathbf{X} = (\mathbf{X}_{[1]}, \ldots, \mathbf{X}_{[k]}, \ldots \mathbf{X}_{[29]})$, using a trivariate Gaussian distribution:

$$\mathbf{X}_{[k]} \sim MVN\left(\begin{pmatrix}0\\0\\0\end{pmatrix}, \begin{pmatrix}1 & \rho & \rho\\ \rho & 1 & \rho\\ \rho & \rho & 1\end{pmatrix}\right).$$

For each gene table, the pairwise correlation among omics $\rho$ was drawn from a $\text{Uniform}(0.2, 0.8)$ distribution. To create a given percentage $p_{\text{aberrant}}$ of aberrant individuals in the population, we selected a number of driver genes $|d_{\text{genes}}|$ as well as a number of driver omics per driver gene $|d_{\text{omics}}|$. Let $i$ represent one of the $n \times p_{\text{aberrant}}$ individuals selected to have an aberrant multi-omic profile, $d_{\text{genes},i}$ the set of randomly selected genes for $i$, and for each of these driver genes $g$, $d_{\text{omics},i,g}$ the set of randomly selected omics for $i$. To create aberrant multi-omic profiles for a selected individual $i$ and gene $g$, the simulated value for $\mathbf{X}_{[g]}[i,]$ (i.e., the $i^{\text{th}}$ row of $\mathbf{X}_{[g]}$) was replaced by a draw from a trivariate Gaussian distribution with covariance matrix as before and mean $\mu_{i,g} = (\mu_1, \mu_2, \mu_3)$, where $\mu_d \sim \text{Uniform}((-6,-4) \cup (4,6))$ if $d \in d_{\text{omics},i,g}$ and 0 otherwise for $d$ = 1, 2, 3.

We evaluated every combination of each of the following parameters: sample size $n$ = {30, 50, 100, 250, 500}, percentage of aberrant individuals $p_{\text{aberrant}}$ = {1%, 5%}, number of driver genes $|d_{\text{genes}}|$ = {1, 2, 3}, and number of driver omics per driver gene $|d_{\text{omics}}|$ = {1, 2}. This corresponds to a total of 5 × 2 × 3 × 2 = 60 settings. For each setting, 200 independent datasets were generated.

We compared the performance of the multi-omic *padma* pathway deviation scores to single-omic padma pathway deviation scores (using the first assay as an arbitrary choice) as well analogous pathway deviation scores calculated using a standard PCA (i.e., where MFA per-table weights were not applied). As a measure of the ability to successfully identify the known aberrant individuals, we calculated the area under the receiver operating characteristic curve (AUC) using multi-omic *padma*, single-omic *padma*, and PCA-based pathway deviation scores. In addition, for multi-omic *padma* we also provide AUC values for detecting the true gene drivers using the per-gene contributions to the individualized pathway deviation scores, averaged over the $n \times p_{\text{aberrant}}$ aberrant individuals; this helps to highlight to what extent the true gene drivers for each aberrant individual are successfully recovered by *padma*.

Results are shown in Supplementary Figures 12-14 and Supplementary Table 6. Overall, we note that when the number of driver genes and/or driver omics increases, it becomes increasingly easy to detect aberrant individuals, regardless of the method. In addition, increasing to $p_{\text{aberrant}}$ = 5% yields slightly lower AUCs than 1% outlier individuals, although AUC values for *padma* and PCA remain high. Within a given combination of $p_{\text{aberrant}}$, $|d_{\text{genes}}|$ and $|d_{\text{omics}}|$, increasing sample sizes lead to slight increases in AUC, and less variable performance across simulations.

Across simulation settings, *padma* and PCA perform very similarly (Supplementary Figures 12-13) for identifying aberrant individuals, particularly for large values of $n$, $|d_{\text{genes}}|$, and $|d_{\text{omics}}|$. However, we do observe a slight advantage for *padma* over the PCA-based approach is observed in settings where $|d_{\text{genes}}| = 2$ for smaller sample sizes (i.e., $n \leq 50$). In the most challenging setting, where $|d_{\text{genes}}| = 2$ and $|d_{\text{omics}}| = 1$, *padma* has slightly higher median and mean AUC values across the 200 simulated datasets for nearly every combination of $n$ and $p_{\text{aberrant}}$ (Supplementary Table 6). In nearly all settings, *padma* single-omics performs the worst, which is unsurprising as only a subset of the data are used and pertinent assays for driver genes may have been omitted. Finally, we remark that multi-omic *padma* has the added advantage of providing a straightforward quantification of the importance of each gene in driving aberrant profiles. The average AUC (across the $n \times p_{\text{aberrant}}$ individuals) for detecting gene drivers shows that driver genes are generally successfully recovered across simulation settings, particularly when multiple omics are perturbed (Supplementary Figure 14).

Taken together, this simulation study illustrates the satisfactory performance of *padma* across a wide range of settings, as well as a slight advantage in identifying aberrant individuals for *padma* compared to a PCA-based alternative, particularly in cases with smaller sample sizes (i.e., $n \leq 50$) and fewer driver genes and omics. We also confirmed that the per-gene contributions to the individualized *padma* pathway deviation scores successfully recover the true gene drivers in all scenarios considered here.

#### References

Abdi, H. and Williams, L.J. (2010) Principal component analysis. *WIREs Computational Statistics*, doi:10.1002/wics.101.

Abdi, H., Williams, L.J., and Valentin, D. (2013) Multiple factor analysis: principal component analysis for multitable and multiblock data sets. *WIREs Computational Statistics*, doi:10.1002/wics.1246.

Akulenko, R., Merl, M. and Helms, V.. (2016). BEclear: Batch Effect Detection and Adjustment in DNA Methylation Data. *PLOS ONE* 11(8), e0159921.

Fabregat, A., Jupe, S., Matthews, L., Sidiropoulos, K., Gillespie, M., Garapati, P., Haw, R., Jassal, B., Korninger, F., May, B., Milacic, M., Roca, C. D., Rothfels, K., Sevilla, C., Shamovsky, V., Shorser, S., Varusai, T., Viteri, G., Weiser, J., Wu, G., Stein, L., Hermjakob, H. and others. (2018). The Reactome Pathway Knowledgebase. *Nucleic Acids Research* 46(D1), D649-D655.

Leek, J. T., Scharpf, R. B., Bravo, H. C., Simcha, D., Langmead, B., Johnson, W. E., Geman, D., Baggerly, K. and Irizarry, R. A. (2010). Tackling the widespread and critical impact of batch effects in high-throughput data. *Nature Reviews Genetics* 11(10), 733-739.

Li, B. and Dewey, C. N. (2011). RSEM: accurate transcript quantification from RNA-Seq data with or without a reference genome. *BMC Bioinformatics* 12(1), 323.

Liberzon, A., Subramanian, A., Pinchback, R., Thorvaldsdottir, H., Tamayo, P. and Mesirov, J. P. (2011). Molecular signatures database (MSigDB) 3.0. *Bioinformatics* 27(12), 1739-1740.

Naba, A., Clauser, K. R., Hoersch, S., Liu, H., Carr, S. A. and Hynes, R. O. (2012). The Matrisome: In Silico Definition and In Vivo Characterization by Proteomics of Normal and Tumor Extracellular Matrices. *Molecular & Cellular Proteomics* 11(4), M111.014647.

Ritchie, M. E., Phipson, B., Wu, D., Hu, Y., Law, C. W., Shi, W. and Smyth, G. K. (2015). Limma powers differential expression analyses for RNA-sequencing and microarray studies. *Nucleic Acids Research* 43(7), e47.

Robinson, M. D. and Oshlack, A. (2010). A scaling normalization method for differential expression analysis of RNA-seq data. *Genome Biology* 11(3), R25.

Schaefer, C. F., Anthony, K., Krupa, S., Buchoff, J., Day, M., Hannay, T. and Buetow, K. H. (2009). PID: the Pathway Interaction Database. *Nucleic Acids Research* 37(suppl 1), D674-D679.

Wan, Y.-W., Allen, G. I. and Liu, Z. (2016). TCGA2stat: simpleTCGA data access for integrated statistical analysis in R. *Bioinformatics* 32(6), 952-954.

### Supplementary Figures

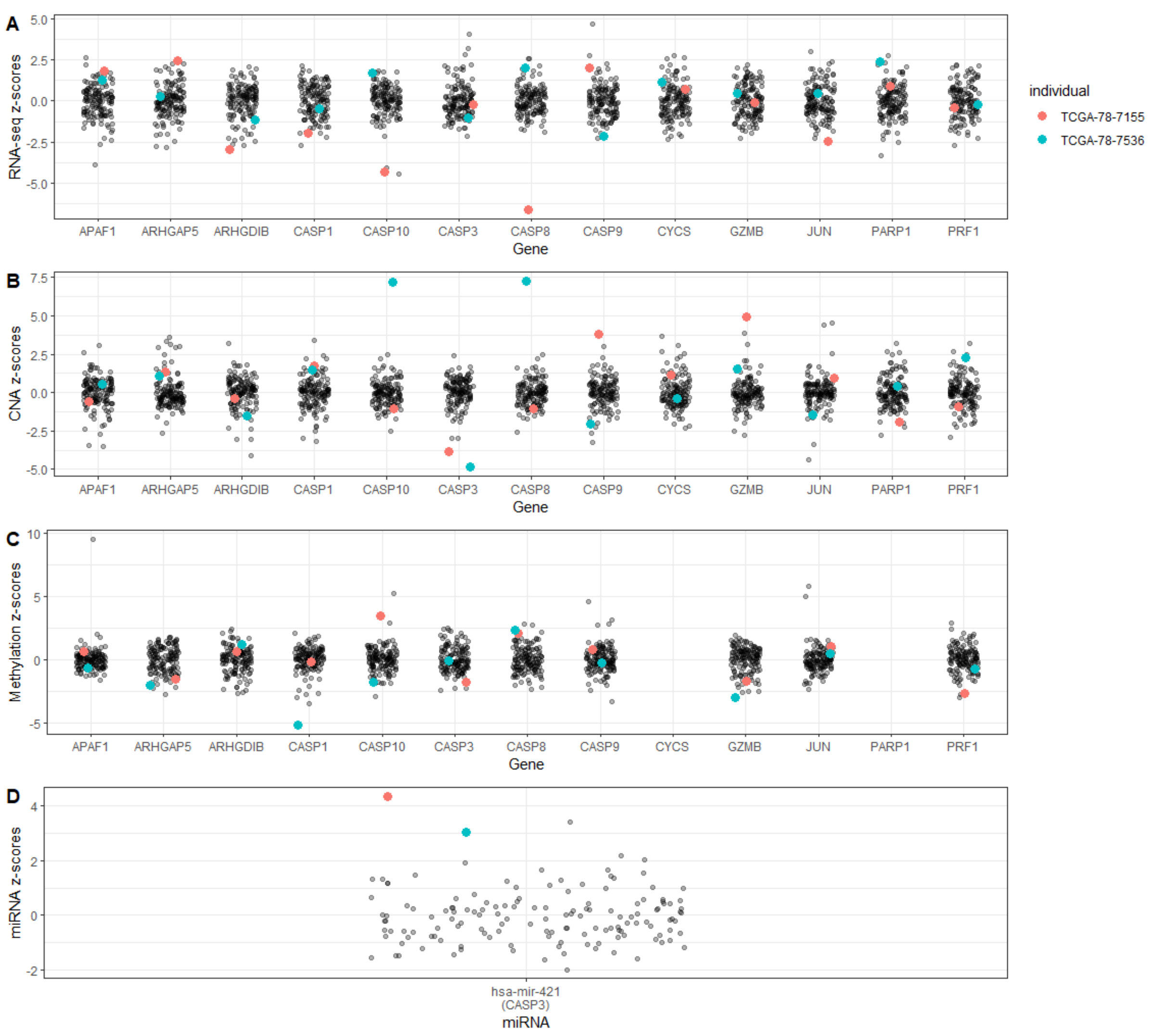

Supplementary Figure 1. Z-scores of RNA-seq, CNA, methylation, and miRNA-seq data for genes in the D4-GDI signaling pathway for individuals in the TCGA LUAD data (n = 144). Data corresponding to the two individuals with the largest overall pathway deviation scores, TCGA-78-7155 and TCGA-78-7536, are highlighted in red and blue.

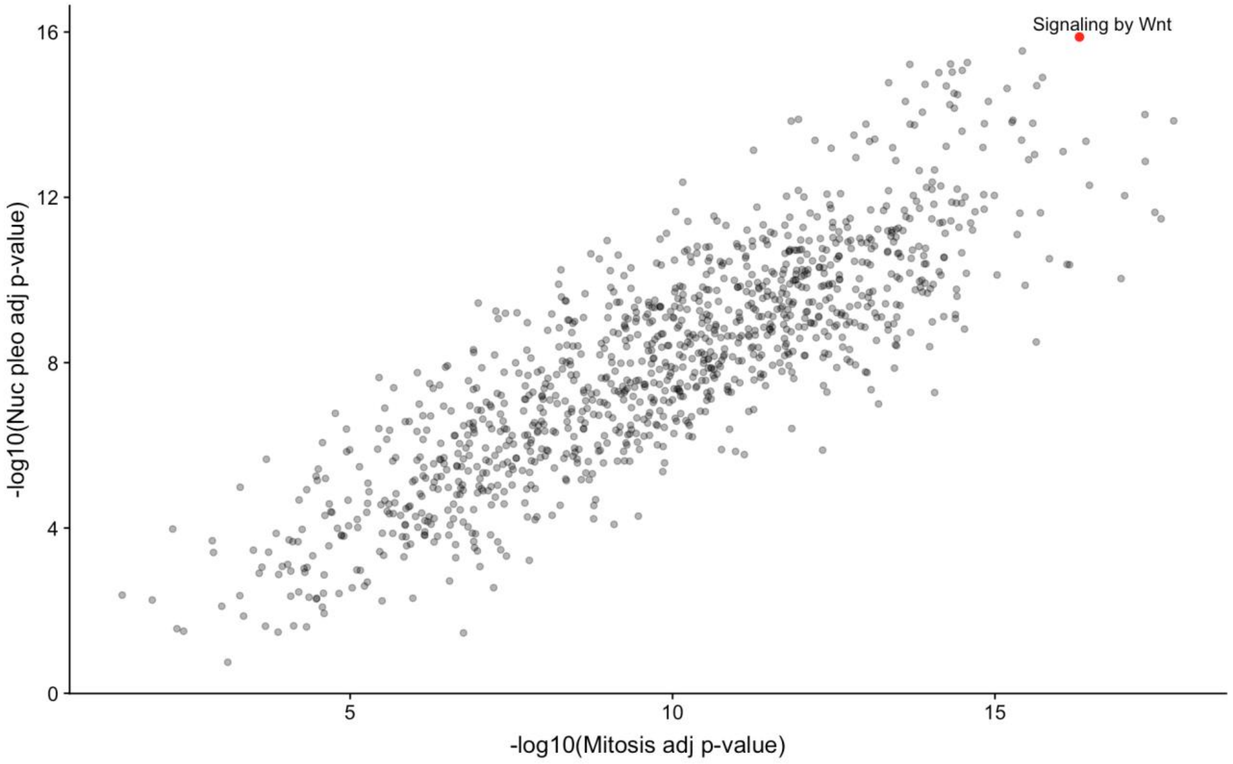

Supplementary Figure 2. Negative log10-transformed p-values from the ANOVA F-test of pathway deviation score versus mitosis and nuclear pleomorphism for each pathway among breast cancer individuals. The signaling by Wnt pathway is highlighted in red.

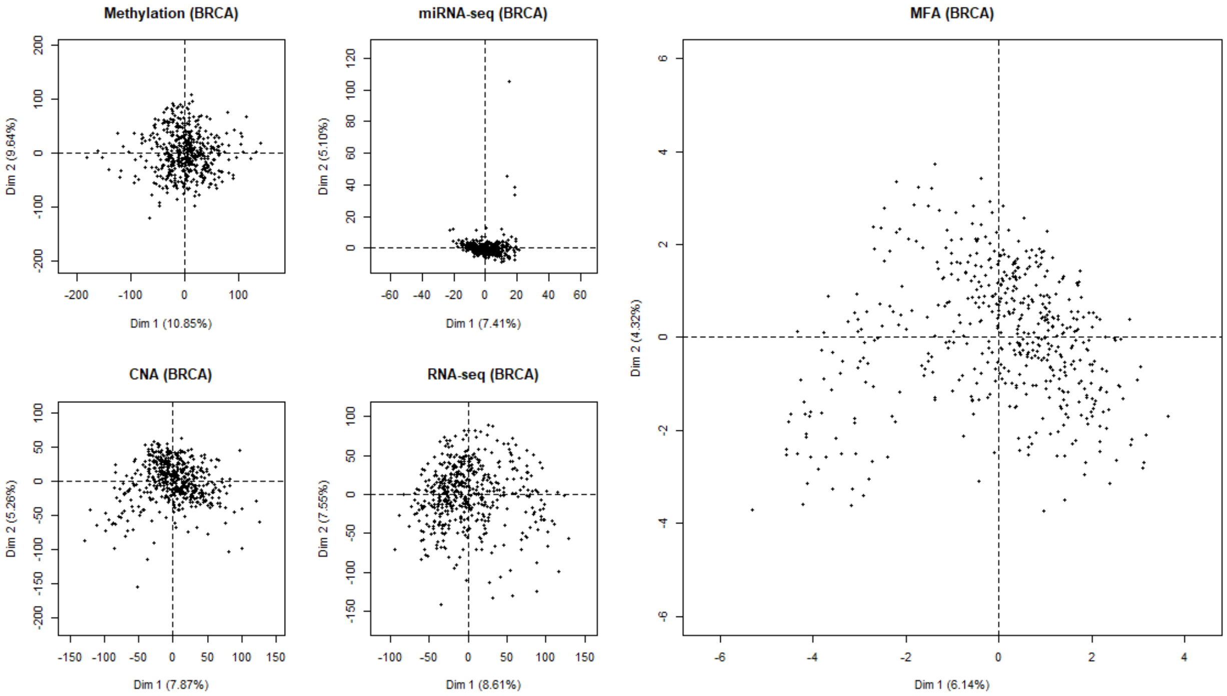

Supplementary Figure 3. Factor maps for the first two dimensions of a global transcriptome- and genome-wide PCA of the methylation, miRNA-seq, CNA, and RNA-seq data (left), as well as a global MFA of all four omics combined (right) for the TCGA BRCA data.

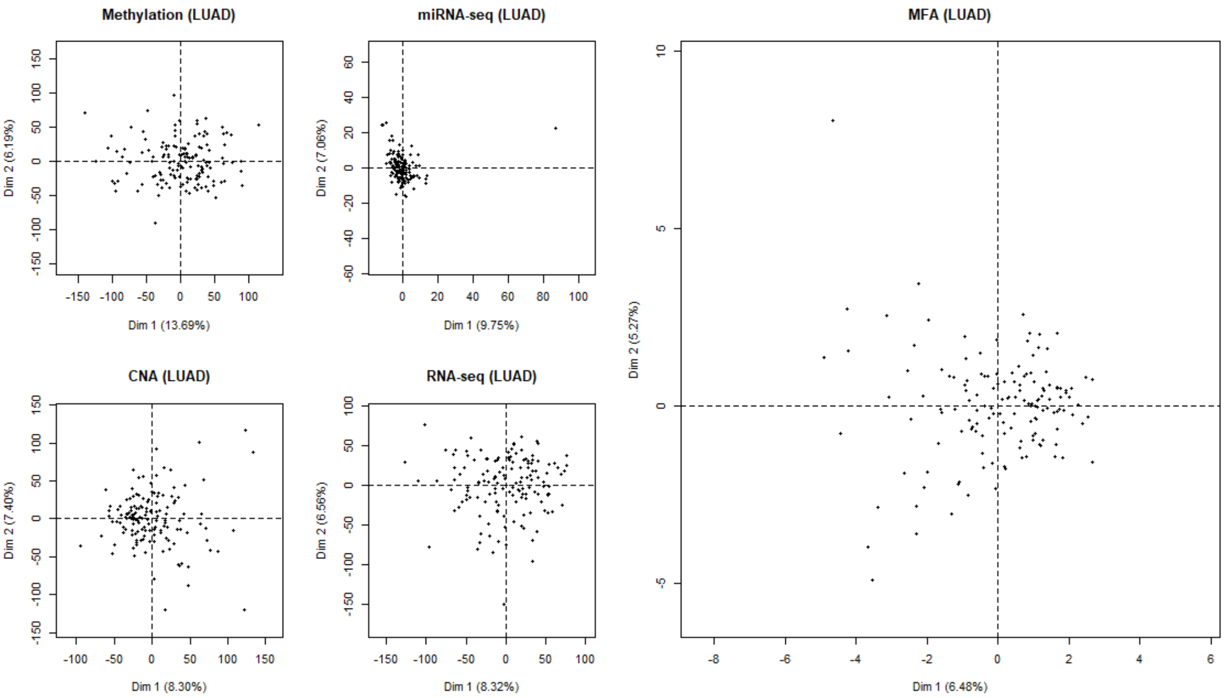

Supplementary Figure 4. Factor maps for the first two dimensions of a global transcriptome- and genome-wide PCA of the methylation, miRNA-seq, CNA, and RNA-seq data (left), as well as a global MFA of all four omics combined (right) for the TCGA LUAD data.

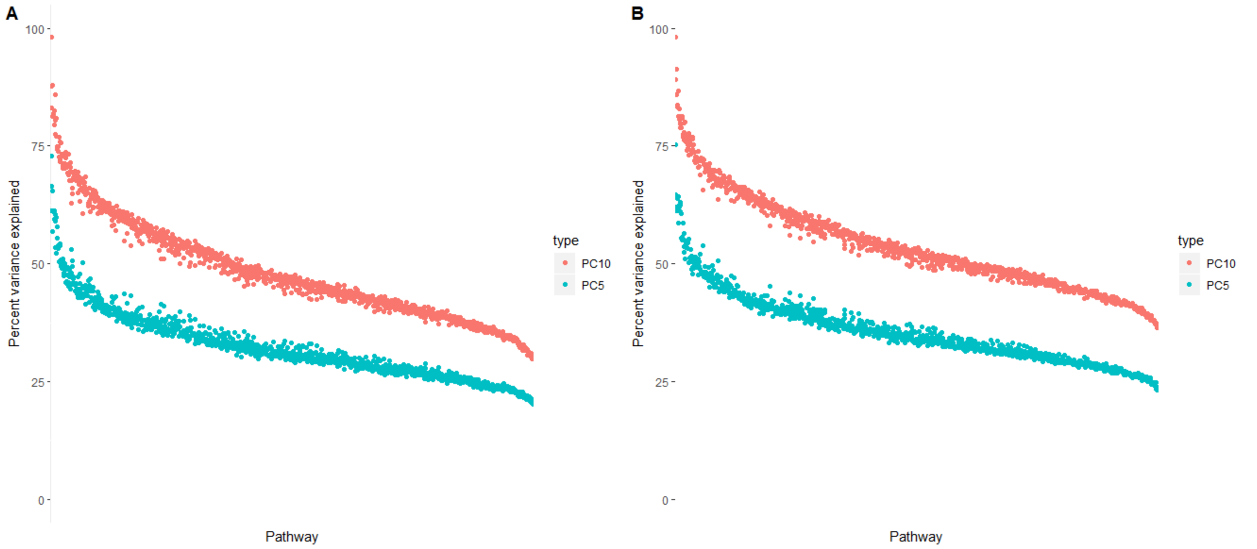

Supplementary Figure 5. Percent variance explained by the first 5 (blue) or 10 (red) components of the MFA for each pathway for the TCGA BRCA (A) and LUAD (B) data.

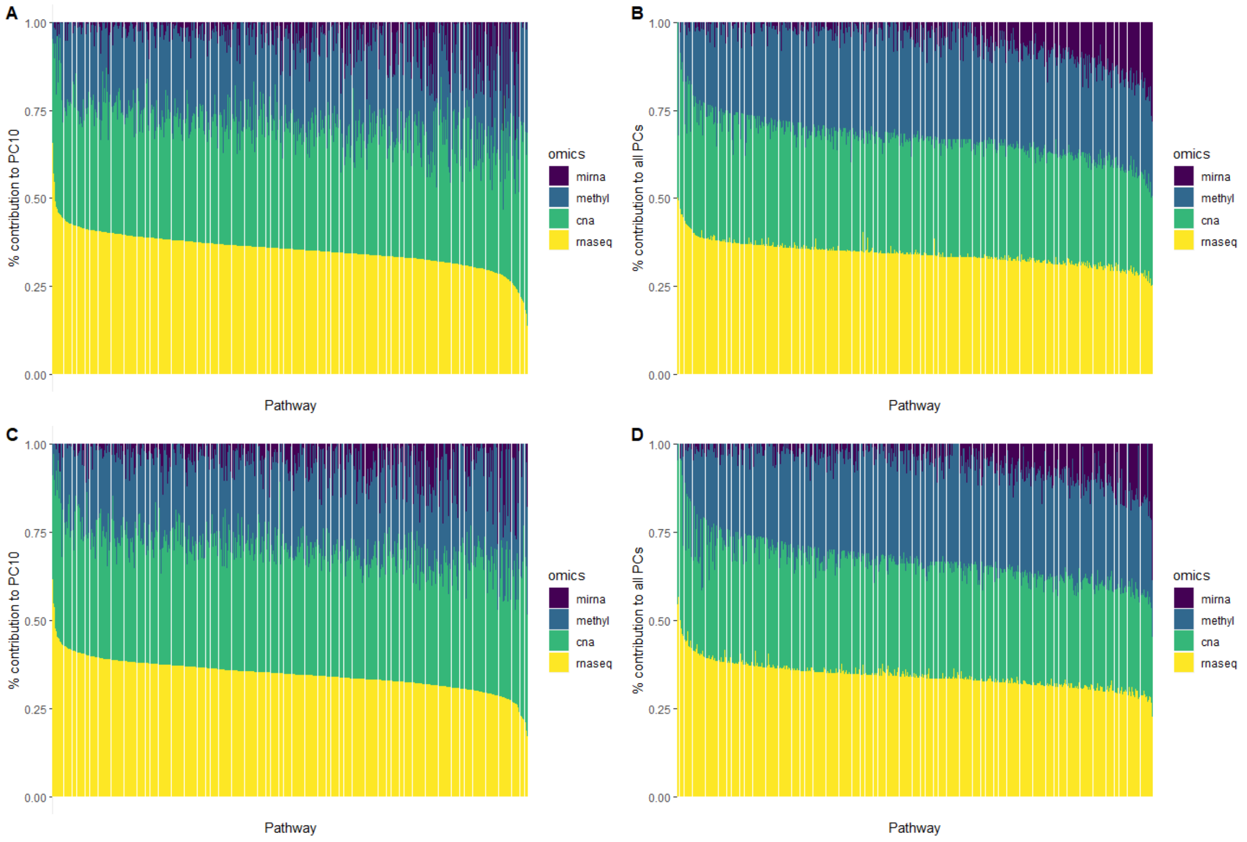

Supplementary Figure 6. Average percent contribution to the MFA of each omic (miRNA-seq, methylation, CNA, RNA-seq) for each pathway. (A) Per-omic average contribution across the first 10 MFA components for TCGA BRCA. (B) Per-omic average contribution across all MFA components for TCGA BRCA. (C) Per-omic average contribution across the first 10 MFA components for TCGA LUAD. (D) Per-omic average contribution across all MFA components for TCGA LUAD.

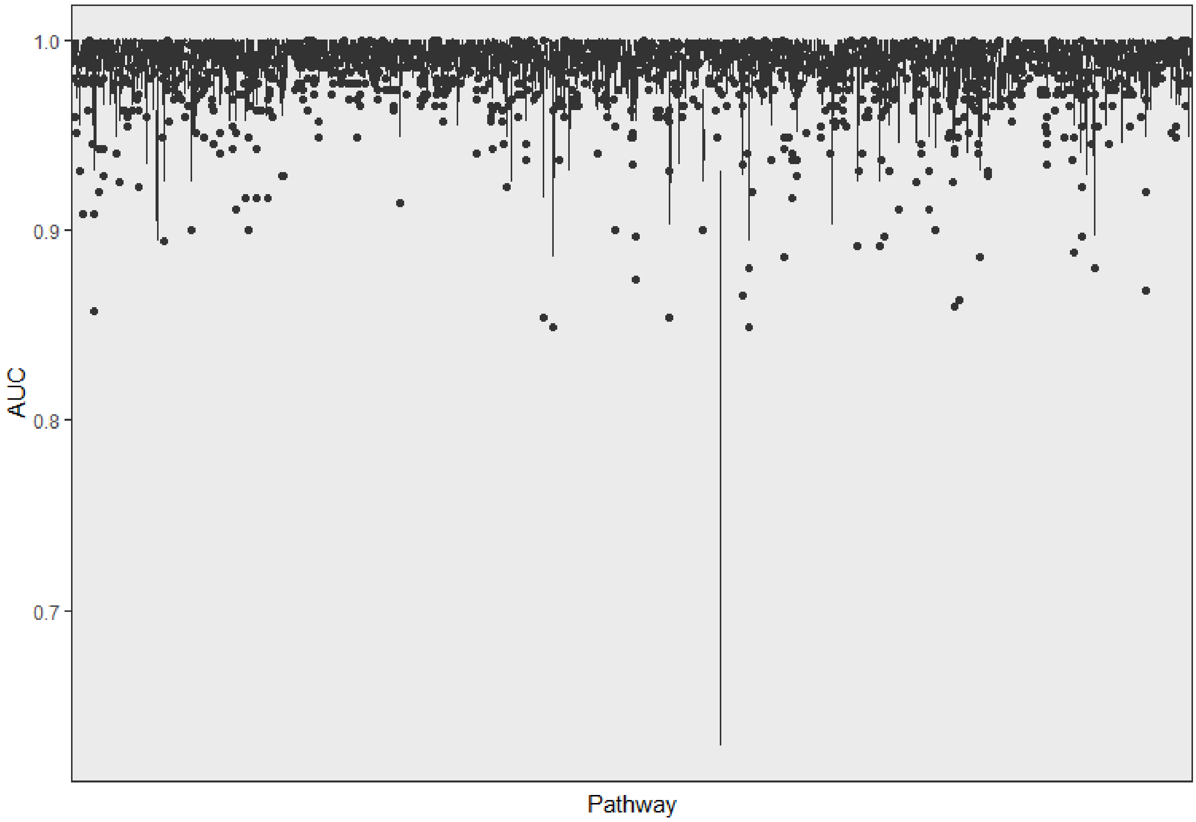

Supplementary Figure 7. Boxplots across 20 independent repetitions of the AUC for detecting 5 randomly selected tumor samples mixed with 70 healthy samples using the *padma* deviation score for the breast cancer multi-omic data (RNA-seq, miRNA-seq, methylation) for each of the 1136 pathways.

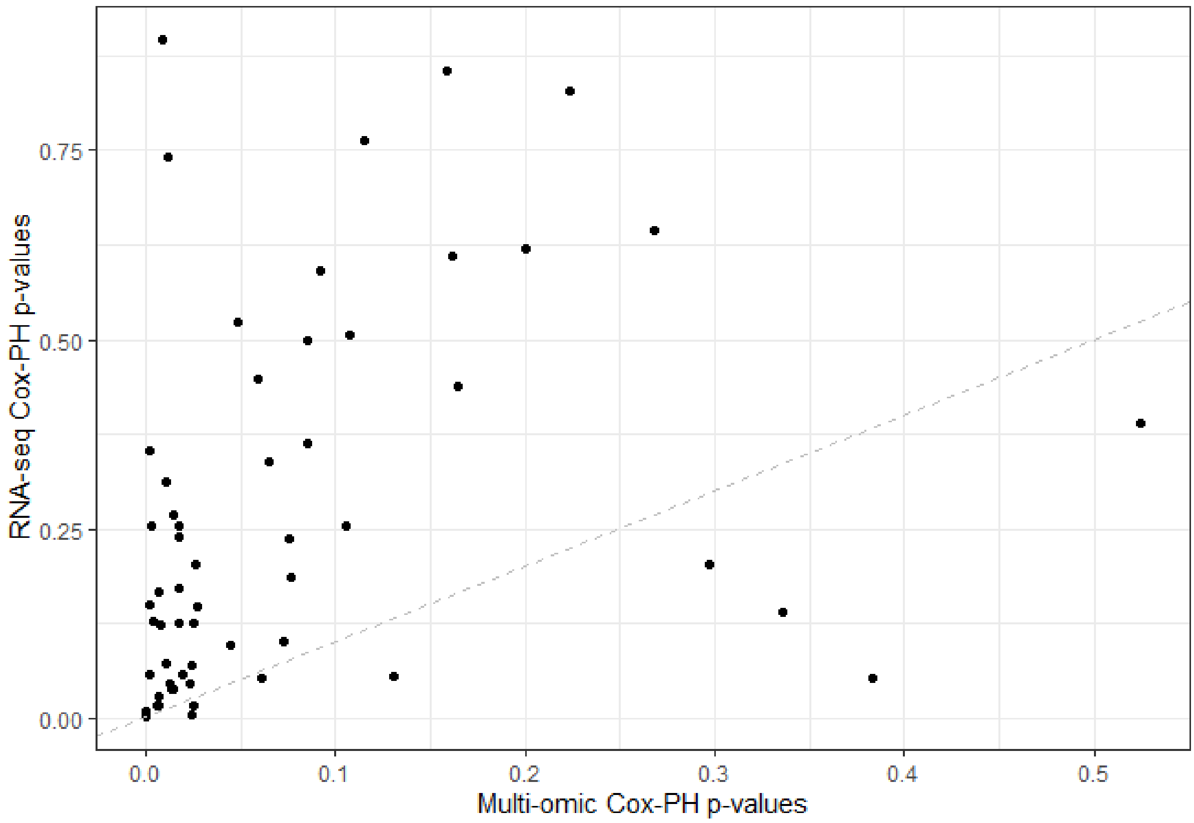

Supplementary Figure 8. Raw p-values from multi-omic and single-omic (RNA-seq only) Cox PH survival analyses in lung cancer for the 57 pathways discussed in the main paper.

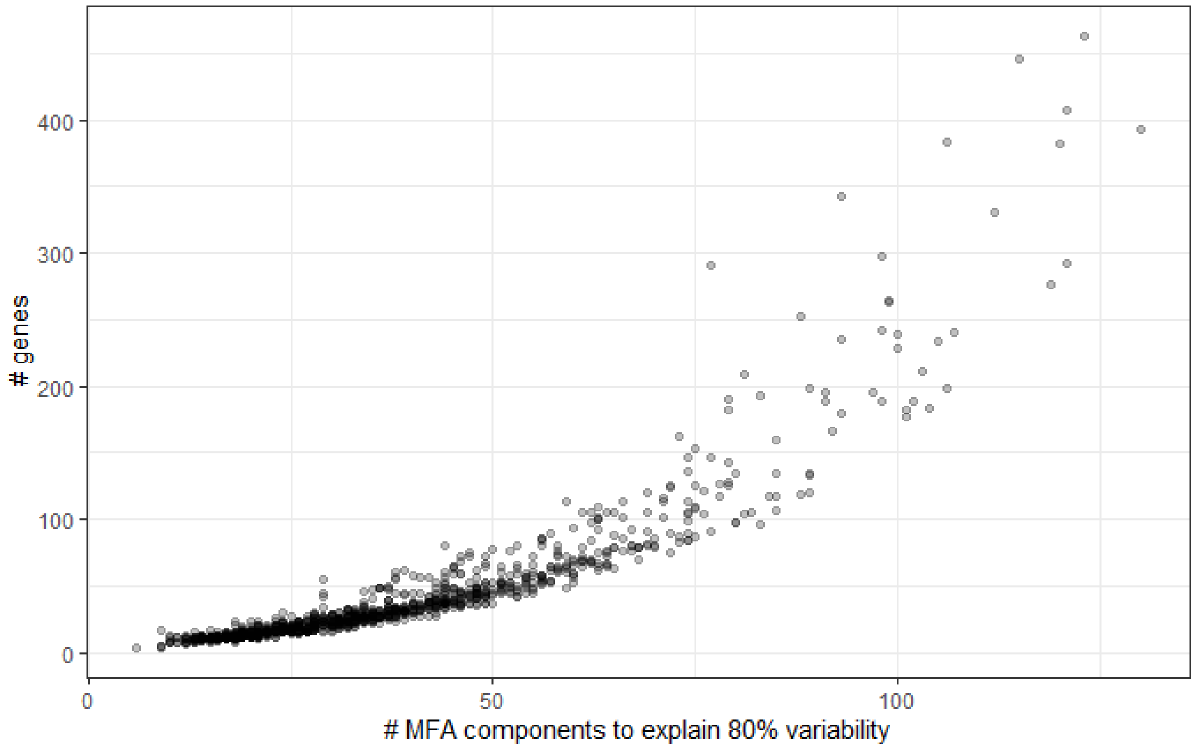

Supplementary Figure 9. Number of MFA components needed to explain 80% of the variability versus the total number of genes for each of the 1136 pathways in the BRCA data studied in the paper.

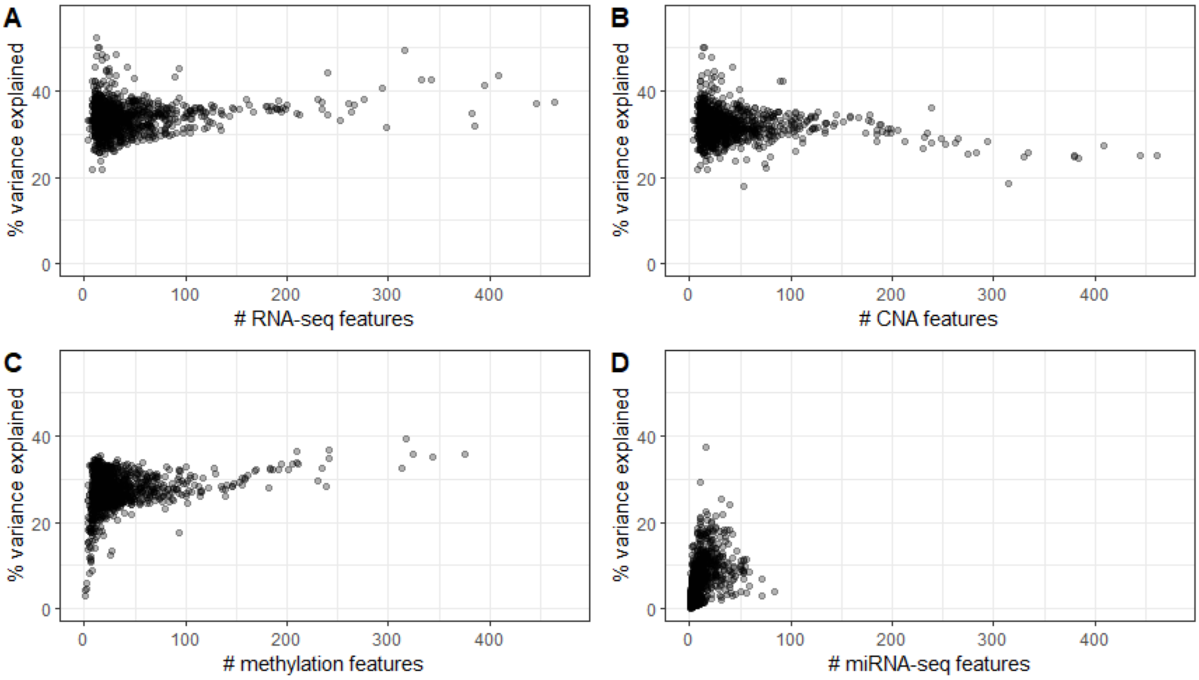

Supplementary Figure 10. Number of available features versus the percent variance explained by the omic across all MFA components for each pathway in the BRCA data.

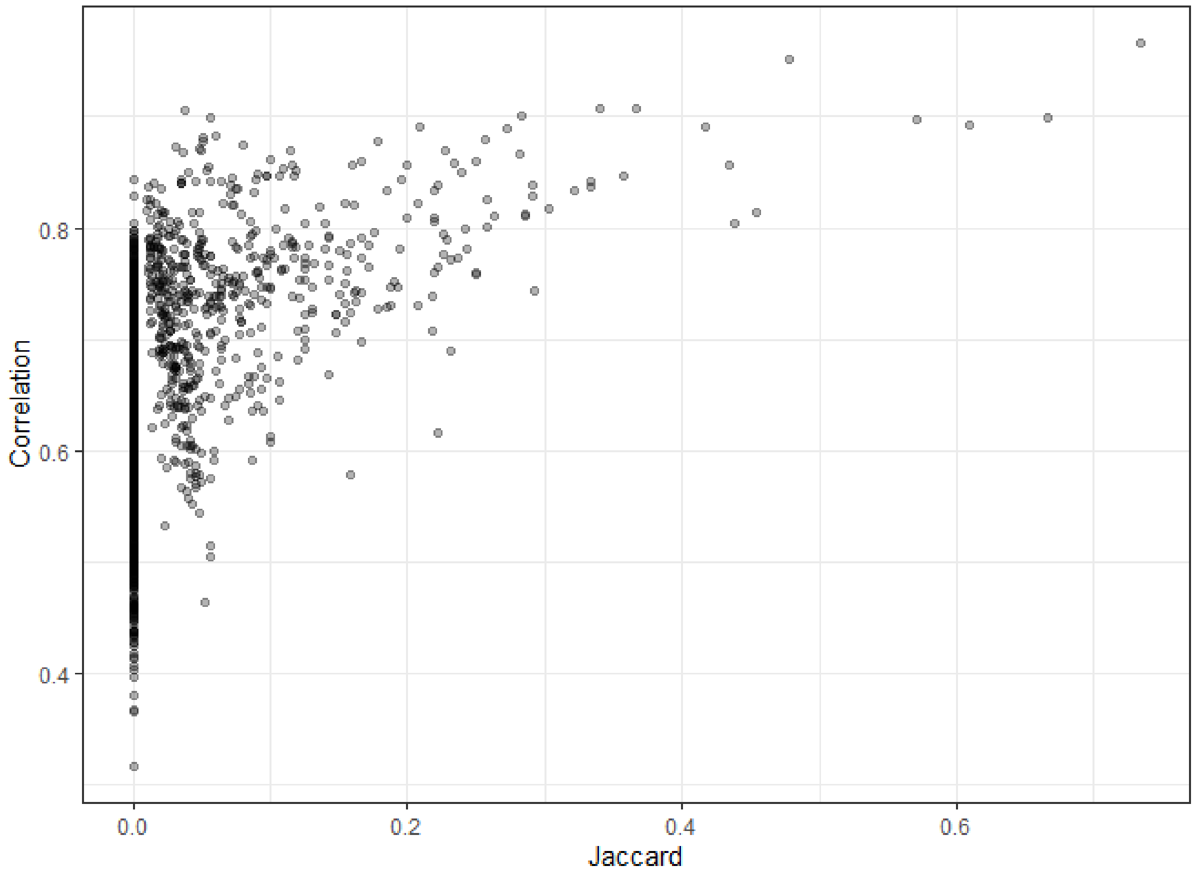

Supplementary Figure 11. Jaccard index versus the Spearman correlation between deviation scores for each pair of pathways among the $n$ = 57 used for the LUAD survival analysis.

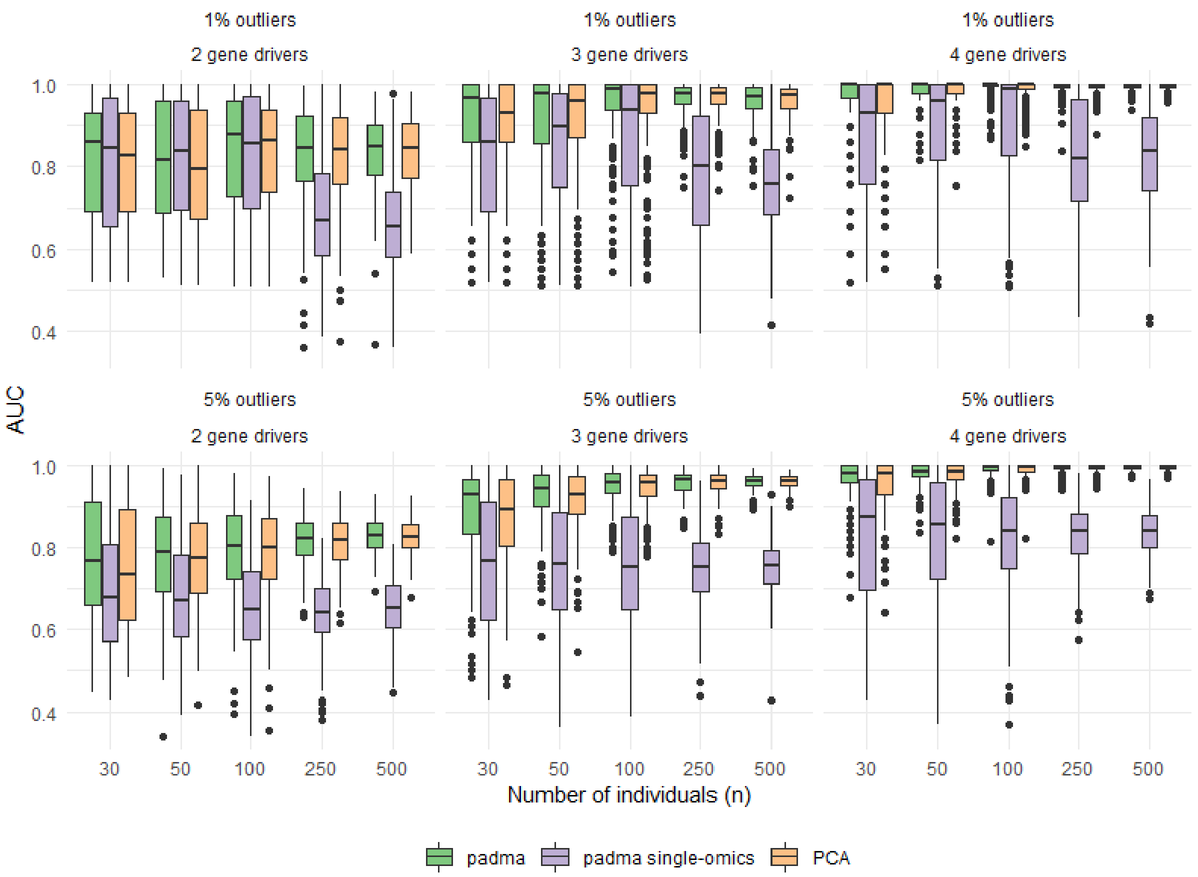

Supplementary Figure 12. Area under the receiver operating characteristic curve (AUC) for identifying aberrant individuals in populations of varying sampling size *n* using individualized pathway deviation scores calculated using multi-omic *padma* (green), single-omics *padma* (purple), and a PCA of concatenated data (orange). Each panel represents a different combination of percentage of outliers in the population ($p_{\text{aberrant}}$ = 1% or 5%) and number of driver genes per aberrant individual ($|d_{\text{genes}}|$ = 2, 3 or 4). Results are shown for $|d_{\text{omics}}|$ = 1 (i.e. a single driver omic per driver gene for each aberrant individual) across 200 simulated datasets.

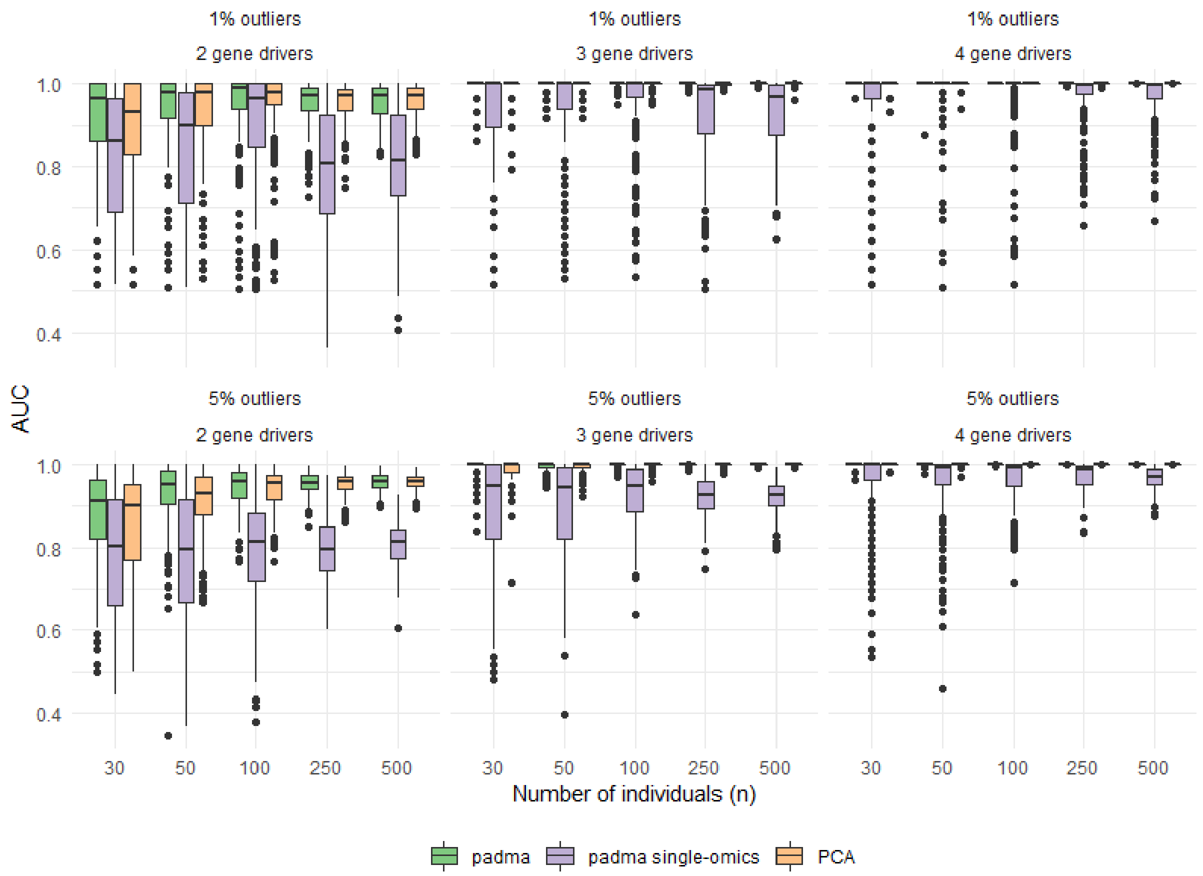

Supplementary Figure 13. Area under the receiver operating characteristic curve (AUC) for identifying aberrant individuals in populations of varying sampling size *n* using individualized pathway deviation scores calculated using multi-omic *padma* (green), single-omics *padma* (purple), and a PCA of concatenated data (orange). Each panel represents a different combination of percentage of outliers in the population ($p_{aberrant}$ = 1% or 5%) and number of driver genes per aberrant individual ($|d_{genes}|$ = 2, 3 or 4). Results are shown for $|d_{omics}|$ = 2 (i.e. two driver omics per driver gene for each aberrant individual) across 200 simulated datasets.

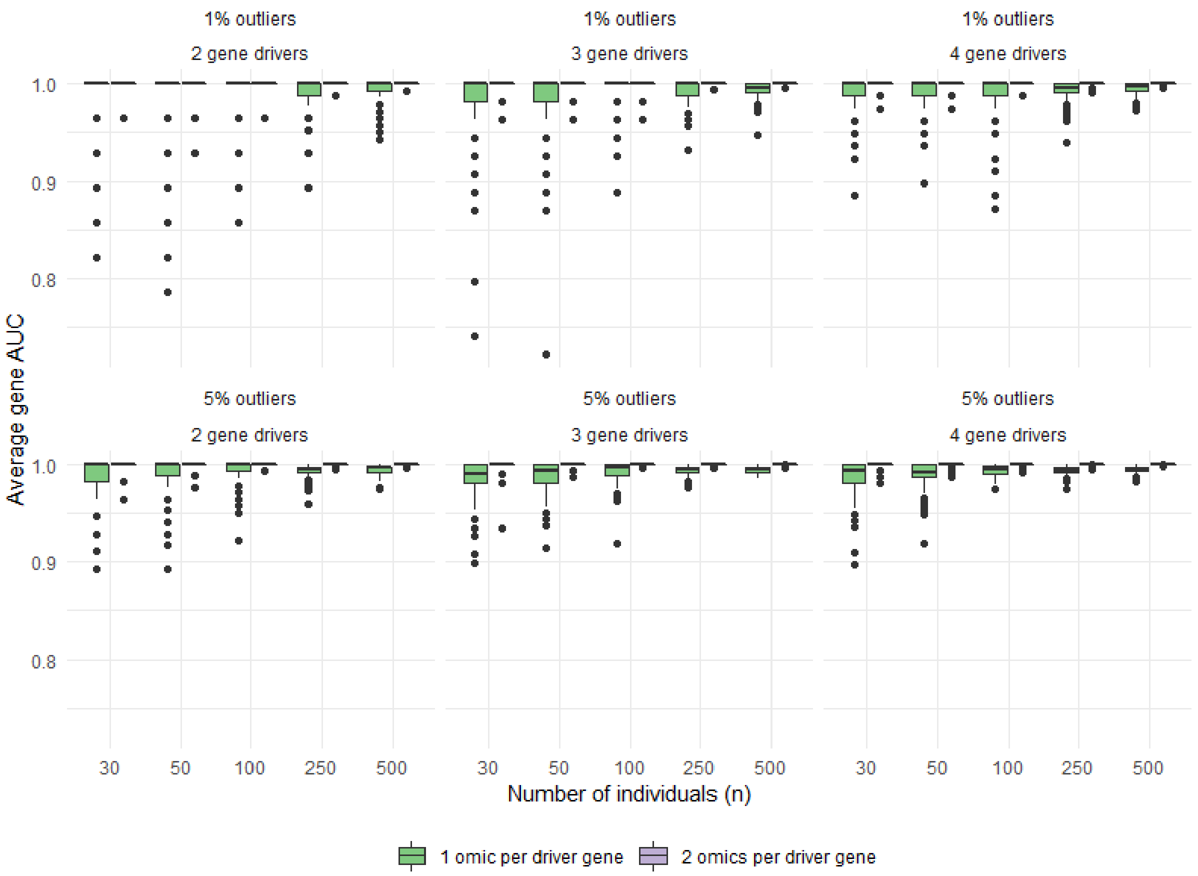

Supplementary Figure 14. Area under the receiver operating characteristic curve (AUC) for identifying the driver genes (averaged across the $p_{\text{aberrant}} \times n$ aberrant individuals in each simulated dataset) using the per-gene contribution to the multi-omic *padma* individual pathway deviation scores in populations of varying sampling size $n$). Each panel represents a different combination of percentage of outliers in the population ($p_{\text{aberrant}}$ = 1% or 5%) and number of driver genes per aberrant individual ($|d_{\text{genes}}|$ = 2, 3 or 4). Results for $|d_{\text{omics}}|$ = 1 and $|d_{\text{omics}}|$ = 2 (i.e., one or two driver omics per driver gene for each aberrant individual) are shown in green and purple, respectively, across 200 simulated datasets.

### Supplementary Tables

Supplementary Table 1. Pathways whose deviation scores are significantly correlated with progression-free interval in lung cancer. Hazard ratios and adjusted p-values correspond to a Cox PH model for pathway deviation, after controlling for age at initial pathologic diagnosis, pathologic tumor stage, and gender, with FDR < 5\%. The number of genes for each pathway corresponds to the number of genes with expression quantified by RNA-seq in the TCGA data.

| Pathway name | Database | Adj. P-value | Hazard ratio | # of genes |
|---|---|---|---|---|
| D4-GDI (GDP dissociation inhibitor) signaling pathway | Biocarta | 0.0155 | 2862 | 13 |
| NF-kB activation through FADD/RIP-1 pathway mediated by caspase-8 and -10 | Reactome | 0.0160 | 1.2996 | 12 |
| CARM1 and regulation of the estrogen receptor | Biocarta | 0.0292 | 1.1619 | 35 |
| Class I PI3K signaling events mediated by AKT | PID | 0.0292 | 1.1761 | 35 |
| Homologous recombination repair of replication-independent double-strand breaks | Reactome | 0.0292 | 1.2798 | 16 |
| ATM signaling pathway | Biocarta | 0.0296 | 1.1692 | 20 |
| G1 and S phases | Sigma-Aldrich | 0.0296 | 1.1859 | 15 |
| CD40L Signaling Pathway | Biocarta | 0.0359 | 1.2165 | 15 |
| p53 signaling pathway | Biocarta | 0.0359 | 1.1752 | 16 |
| Hypoxia and p53 in the Cardiovascular system | Biocarta | 0.0359 | 1.1911 | 22 |
| Double stranded RNA induced gene expression | Biocarta | 0.0359 | 1.2301 | 10 |
| Regulation of telomerase | PID | 0.0359 | 1.1075 | 68 |
| Binding and entry of HIV virion | Reactome | 0.0379 | 1.3202 | 4 |
| Cell cycle: G2/M checkpoint | Biocarta | 0.0409 | 1.1646 | 24 |
| Sumoylation by RanBP2 regulates transcriptional repression | PID | 0.0409 | 1.2244 | 11 |
| Role of BRCA1, BRCA2 and ATR in Cancer Susceptibility | Biocarta | 0.0422 | 1.1856 | 21 |
| Cell Cycle: G1/S Check Point | Biocarta | 0.0422 | 1.1295 | 28 |
| Role of ERBB2 in Signal Transduction and Oncology | Biocarta | 0.0422 | 1.1592 | 22 |
| Influence of Ras and Rho proteins on G1 to S Transition | Biocarta | 0.0422 | 1.1220 | 26 |

| | | | | |
|---|---|---|---|---|
| Sprouty regulation of tyrosine kinase signals | Biocarta | 0.0422 | 1.1822 | 18 |
| Signaling events mediated by HDAC Class III | PID | 0.0422 | 1.1582 | 25 |
| Fanconi Anemia Pathway | Reactome | 0.0422 | 1.1848 | 21 |
| Glycolysis | Reactome | 0.0422 | 1.1582 | 27 |
| TNF-type receptor Fas induces apoptosis on ligand binding | Sigma-Aldrich | 0.0422 | 1.2082 | 9 |
| AKT phosphorylates targets in the cytosol | Reactome | 0.0454 | 1.1722 | 12 |
| Induction of apoptosis through DR3 and DR4/5 Death Receptors | Biocarta | 0.0484 | 1.1152 | 33 |
| Erythropoietin mediated neuroprotection through NF-kB | Biocarta | 0.0484 | 1.1769 | 11 |
| Telomeres, Telomerase, Cellular Aging, and Immortality | Biocarta | 0.0484 | 1.1239 | 18 |
| a6b1 and a6b4 Integrin signaling | PID | 0.0484 | 1.0994 | 46 |
| HIV-1 Nef: Negative effector of Fas and TNF-alpha | PID | 0.0484 | 1.1246 | 35 |
| GAB1 signalosome | Reactome | 0.0484 | 1.1216 | 36 |
| SHC1 events in EGFR signaling | Reactome | 0.0484 | 1.1586 | 15 |

Supplementary Table 2. Pathways whose deviation scores are significantly correlated with measures of histological grade (mitosis, nuclear pleomorphism) in breast cancer. Adjusted p-values after Benjamini-Hochberg correction were $< 3.31 \times 10^{-12}$ for all pathways presented in the table. Combined ranks correspond to the rank product of the individual rankings from mitosis and nuclear pleomorphism, and the number of genes for each pathway corresponds to the number of genes with expression quantified by RNA-seq in the TCGA data.

| **Pathway name** | **Database** | **Combined ranking** | **# of genes** |
|---|---|---|---|
| Signaling by Wnt | Reactome | 3.16 | 63 |
| Apoptotic execution phase | Reactome | 5.00 | 52 |
| APC/C:Cdh1 mediated degradation of Cdc20 and other APC/C:Cdh1 targeted proteins in late mitosis/early G1 | Reactome | 6.78 | 64 |
| Genes involved in Beta-catenin phosphorylation cascade | Reactome | 10.49 | 16 |
| Autodegradation of Cdh1 by Cdh1:APC/C | Reactome | 10.95 | 56 |
| Genes involved in M/G1 transition | Reactome | 11.62 | 72 |
| Regulation of the Fanconi anemia pathway | Reactome | 13.93 | 7 |
| Apoptotic cleavage of cellular proteins | Reactome | 14.14 | 38 |
| Apoptosis | Reactome | 14.28 | 143 |
| ER-phagosome pathway | Reactome | 15.62 | 58 |

Supplementary Table 3. Sample size for each histological measure for the *n* = 504 breast cancer patients.

| **Histological measure** | **Stage** | **Number of individuals (*n* = 504)** |
|---|---|---|
| Nuclear pleomorphism | I: Small regular nuclei<br>II: Moderate increase in size<br>III: Moderate to marked variation in size<br>NA | 47<br>207<br>162<br>87 |
| Mitotic index | I: 0-5 per 10 high powered fields (HPF; low)<br>II: 6-10 per 10 HPF (medium)<br>III: >10 per 10 HPF (high)<br>NA | 225<br>89<br>101<br>89 |
| Glandular/tubule formation | I: >75% (well differentiated)<br>II: 10-75% (moderately differentiated)<br>III: < 10% (poorly differentiated)<br>NA | 51<br>81<br>285<br>87 |

Supplementary Table 4. Full gene lists for pathways in Supplementary Table 1. Genes correspond to those with expression quantified by RNA-seq in the TCGA data.

| **Pathway name** | **Genes** |
|---|---|
| D4-GDI (GDP dissociation inhibitor) signaling pathway | APAF1, ARHGAP5, ARHGDIB, CASP1, CASP10, CASP3, CASP8, CASP9, CYCS, GZMB, JUN, PARP1, PRF1 |
| NF-kB activation through FADD/RIP-1 pathway mediated by caspase-8 and -10 | CASP10, CASP8, CHUK, DDX58, FADD, IFIH1, IKBKB, IKBKG, MAVS, RIPK1, RNF135, TRIM25 |
| CARM1 and regulation of the estrogen receptor | BRCA1, CARM1, CCND1, CREBBP, EP300, ERCC3, ESR1, GRIP1, GTF2A1, GTF2E1, GTF2F1, HDAC1, HDAC10, HDAC11, HDAC2, HDAC3, HDAC4, HDAC5, HDAC6, HDAC7, HDAC8, HDAC9, HIST2H3C, MED1, MEF2C, NCOR2, NR0B1, NRIP1, PELP1, PHB2, POLR2A, PPARGC1A, SPEN, SRA1, TBP |
| Class I PI3K signaling events mediated by AKT | AKT1, AKT2, AKT3, BAD, BCL2L1, CASP9, CDKN1A, CDKN1B, CHUK, FOXO1, FOXO3, FOXO4, GSK3A, GSK3B, HSP90AA1, KPNA1, MAP3K5, MAPKAP1, MLST8, MTOR, PDPK1, PRKACA, PRKDC, RAF1, RICTOR, SFN, SLC2A4, SRC, TBC1D4, YWHAB, YWHAE, YWHAG, YWHAH, YWHAQ, YWHAZ |
| Homologous recombination repair of replication-independent double-strand breaks | ATM, BRCA1, BRCA2, BRIP1, H2AFX, LIG1, MDC1, MRE11A, NBN, RAD50, RAD51, RAD52, RPA1, RPA2, RPA3, TP53BP1 |
| ATM signaling pathway | ABL1, ATM, BRCA1, CDKN1A, CHEK1, CHEK2, GADD45A, JUN, MAPK8, MDM2, MRE11A, NBN, NFKB1, NFKBIA, RAD50, RAD51, RBBP8, RELA, TP53, TP73 |
| G1 and S phases | ARF1, ARF3, CCND1, CDK2, CDK4, CDKN1A, CDKN1B, CDKN2A, CFL1, E2F1, E2F2, MDM2, NXT1, PRB1, TP53 |
| CD40L Signaling Pathway | CD40, CD40LG, CHUK, DUSP1, IKBKAP, IKBKB, IKBKG, MAP3K1, MAP3K14, NFKB1, NFKBIA, RELA, TNFAIP3, TRAF3, TRAF6 |
| p53 signaling pathway | APAF1, ATM, BAX, BCL2, CCND1, CCNE1, CDK2, CDK4, CDKN1A, E2F1, GADD45A, MDM2, PCNA, RB1, TIMP3, TP53 |
| Hypoxia and p53 in the Cardiovascular system | ABCB1, AKT1, ATM, BAX, CDKN1A, CSNK1A1, CSNK1D, EP300, FHL2, GADD45A, HIC1, HIF1A, HSP90AA1, HSPA1A, IGFBP3, MAPK8, MDM2, NFKBIB, NQO1, RPA1, TAF1, TP53 |
| Double stranded RNA induced gene expression | CHUK, DNAJC3, EIF2AK2, EIF2S1, EIF2S2, MAP3K14, NFKB1, NFKBIA, RELA, TP53 |
| Regulation of telomerase | ABL1, ACD, AKT1, ATM, BLM, CCND1, CDKN1B, DKC1, E2F1, EGF, EGFR, ESR1, FOS, HDAC1, HDAC2, HNRNPC, HSP90AA1, HUS1, IFNAR2, IFNG, IL2, IRF1, JUN, MAPK1, MAPK3, MAX, MRE11A, MTOR, MXD1, MYC, NBN, NCL, NFKB1, NR2F2, PARP2, PIF1, PINX1, POT1, PTGES3, RAD1, RAD50, RAD9A, RBBP4, RBBP7, RPS6KB1, SAP18, SAP30, SIN3A, SIN3B, SMAD3, SMG5, SMG6, SP1, SP3, TERF1, TERF2, TERF2IP, TERT, TGFB1, TINF2, TNKS, UBE3A, WRN, WT1, XRCC5, XRCC6, YWHAE, ZNFX1 |
| Binding and entry of HIV virion | CCR5, CD4, CXCR4, PPIA |
| Cell cycle: G2/M checkpoint | ATM, ATR, BRCA1, CCNB1, CDC25A, CDC25B, CDC25C, CDC34, CDK1, CDKN1A, CDKN2D, CHEK1, CHEK2, EP300, GADD45A, MDM2, MYT1, PLK1, PRKDC, RPS6KA1, TP53, WEE1, YWHAH, YWHAQ |
| Sumoylation by RanBP2 regulates transcriptional repression | HDAC1, HDAC4, MDM2, PIAS1, PIAS2, RAN, RANBP2, RANGAP1, SUMO1, UBE2I, XPO1 |
| Role of BRCA1, BRCA2 and ATR in Cancer Susceptibility | ATM, ATR, BRCA1, BRCA2, CHEK1, CHEK2, FANCC, FANCD2, FANCE, FANCF, FANCG, HUS1, MRE11A, NBN, RAD1, RAD17, RAD50, RAD51, RAD9A, TP53, TREX1 |

| | |
|---|---|
| Cell Cycle: G1/S Check Point | ABL1, ATM, ATR, CCNA1, CCND1, CCNE1, CDC25A, CDK1, CDK2, CDK4, CDK6, CDKN1A, CDKN1B, CDKN2A, CDKN2B, DHFR, E2F1, GSK3B, HDAC1, RB1, SKP2, SMAD3, SMAD4, TFDP1, TGFB1, TGFB2, TGFB3, TP53 |
| Role of ERBB2 in Signal Transduction and Oncology | CARM1, EGFR, EP300, ERBB3, ERBB4, ESR1, GRB2, GRIP1, HRAS, IL6, IL6R, IL6ST, MAP2K1, MAPK1, MAPK3, PIK3CA, PIK3CG, PIK3R1, RAF1, SHC1, SOS1, STAT3 |
| Influence of Ras and Rho proteins on G1 to S Transition | AKT1, CCND1, CCNE1, CDK2, CDK4, CDK6, CDKN1A, CDKN1B, CHUK, E2F1, HRAS, IKBKB, IKBKG, MAPK1, MAPK3, NFKB1, NFKBIA, PAK1, PIK3CA, PIK3R1, RAC1, RAF1, RB1, RELA, RHOA, TFDP1 |
| Sprouty regulation of tyrosine kinase signals | CBL, EGF, EGFR, GRB2, HRAS, MAP2K1, MAPK1, MAPK3, PTPRB, RAF1, RASA1, SHC1, SOS1, SPRY1, SPRY2, SPRY3, SPRY4, SRC |
| Signaling events mediated by HDAC Class III | ACSS1, ACSS2, BAX, CDKN1A, CREBBP, EP300, FHL2, FOXO1, FOXO3, FOXO4, HDAC4, HIST1H1E, HOXA10, KAT2B, MEF2D, MYOD1, PPARGC1A, SIRT1, SIRT2, SIRT3, SIRT7, TP53, TUBA1B, TUBB2A, XRCC6 |
| Fanconi Anemia Pathway | ATM, ATR, BRCA1, BRCA2, C17orf70, C19orf40, FANCA, FANCB, FANCC, FANCD2, FANCE, FANCF, FANCG, FANCL, FANCM, PALB2, RPS27A, UBA52, UBE2T, USP1, ZBTB32 |
| Glycolysis | ALDOA, ALDOB, ALDOC, ENO1, ENO2, ENO3, GAPDH, GAPDHS, GPI, PFKFB1, PFKFB2, PFKFB3, PFKFB4, PFKL, PFKM, PFKP, PGAM1, PGAM2, PGK1, PKLR, PKM2, PPP2CA, PPP2CB, PPP2R1A, PPP2R1B, PPP2R5D, TPI1 |
| TNF-type receptor Fas induces apoptosis on ligand binding | BCL2, CASP3, CASP8, CFL1, CFLAR, ENDOU, FAS, FASLG, PDE6D |
| AKT phosphorylates targets in the cytosol | AKT1, AKT1S1, AKT2, AKT3, BAD, CASP9, CDKN1A, CDKN1B, CHUK, GSK3A, MDM2, TSC2 |
| Induction of apoptosis through DR3 and DR4/5 Death Receptors | APAF1, BCL2, BID, BIRC2, BIRC3, CASP10, CASP3, CASP6, CASP7, CASP8, CASP9, CFLAR, CHUK, CYCS, DFFA, DFFB, FADD, GAS2, LMNA, MAP3K14, NFKB1, NFKBIA, RELA, RIPK1, SPTAN1, TNFRSF10A, TNFRSF10B, TNFRSF25, TNFSF10, TNFSF12, TRADD, TRAF2, XIAP |
| Erythropoietin mediated neuroprotection through NF-kB | ARNT, CDKN1A, EPO, EPOR, GRIN1, HIF1A, JAK2, NFKB1, NFKBIA, RELA, SOD2 |
| Telomeres, Telomerase, Cellular Aging, and Immortality | AKT1, BCL2, EGFR, HSP90AA1, IGF1R, KRAS, MYC, POLR2A, PPP2CA, PRKCA, RB1, TEP1, TERF1, TERT, TNKS, TP53, XRCC5, XRCC6 |
| a6b1 and a6b4 Integrin signaling | AKT1, CASP7, CD9, CDH1, COL17A1, EGF, EGFR, ERBB2, ERBB3, GRB2, HRAS, IL1A, ITGA6, ITGB1, ITGB4, LAMA1, LAMA2, LAMA3, LAMA4, LAMA5, LAMB1, LAMB2, LAMB3, LAMC1, LAMC2, LAMC3, MET, MST1, MST1R, PIK3CA, PIK3R1, PMP22, PRKCA, RAC1, RPS6KB1, RXRA, RXRB, RXRG, SFN, SHC1, YWHAB, YWHAE, YWHAG, YWHAH, YWHAQ, YWHAZ |
| HIV-1 Nef: Negative effector of Fas and TNF-alpha | APAF1, BAG4, BCL2, BID, BIRC3, CASP2, CASP3, CASP6, CASP7, CASP8, CASP9, CD247, CFLAR, CHUK, CRADD, CYCS, DAXX, DFFA, DFFB, FADD, FAS, FASLG, MAP2K7, MAP3K14, MAP3K5, MAPK8, NFKB1, NFKBIA, RELA, RIPK1, TNF, TNFRSF1A, TRADD, TRAF1, TRAF2 |
| GAB1 signalosome | AKT1, AKT1S1, AKT2, AKT3, BAD, CASP9, CDKN1A, CDKN1B, CHUK, CREB1, CSK, EGF, EGFR, FOXO1, FOXO3, FOXO4, GAB1, GRB2, GSK3A, MAPKAP1, MDM2, MLST8, MTOR, NR4A1, PAG1, PDPK1, PHLPP1, PIK3CA, PIK3R1, PTEN, RICTOR, RPS6KB2, SRC, THEM4, TRIB3, TSC2 |
| SHC1 events in EGFR signaling | CDK1, EGF, EGFR, GRB2, HRAS, KRAS, MAP2K1, MAP2K2, MAPK1, MAPK3, NRAS, RAF1, SHC1, SOS1, YWHAB |

Supplementary Table 5. Full gene lists for pathways in Supplementary Table 2. Genes correspond to those with expression quantified by RNA-seq in the TCGA data.

| **Pathway name** | **Genes** |
|---|---|
| Signaling by Wnt | APC, AXIN1, BTRC, CSNK1A1, CTNNB1, CUL1, FAM123B, FRAT1, FRAT2, PPP2CA, PPP2CB, PPP2R1A, PPP2R1B, PPP2R5A, PPP2R5B, PPP2R5C, PPP2R5D, PPP2R5E, PSMA1, PSMA2, PSMA3, PSMA4, PSMA5, PSMA6, PSMA7, PSMA8, PSMB1, PSMB10, PSMB2, PSMB3, PSMB4, PSMB5, PSMB6, PSMB7, PSMB8, PSMB9, PSMC1, PSMC2, PSMC3, PSMC4, PSMC5, PSMC6, PSMD1, PSMD10, PSMD11, PSMD12, PSMD13, PSMD14, PSMD2, PSMD3, PSMD4, PSMD5, PSMD6, PSMD7, PSMD8, PSMD9, PSME1, PSME2, PSME4, PSMF1, RPS27A, SKP1, UBA52 |
| Apoptotic execution phase | ACIN1, ADD1, APC, BCAP31, BIRC2, BMX, CASP3, CASP6, CASP7, CASP8, CDH1, CTNNB1, DBNL, DFFA, DFFB, DNM1L, DSG1, DSG2, DSG3, DSP, FNTA, GAS2, GSN, H1F0, HIST1H1A, HIST1H1B, HIST1H1C, HIST1H1D, HIST1H1E, HMGB1, HMGB2, KPNA1, KPNB1, LMNA, LMNB1, LOC647859, MAPT, MST4, OCLN, PAK2, PKP1, PLEC, PRKCD, PRKCQ, PTK2, ROCK1, SATB1, SPTAN1, STK24, TJP1, TJP2, VIM |
| APC/C:Cdh1 mediated degradation of Cdc20 and other APC/C:Cdh1 targeted proteins in late mitosis/early G1 | ANAPC1, ANAPC10, ANAPC11, ANAPC2, ANAPC4, ANAPC5, ANAPC7, AURKA, AURKB, CDC16, CDC20, CDC23, CDC26, CDC27, PLK1, PSMA1, PSMA2, PSMA3, PSMA4, PSMA5, PSMA6, PSMA7, PSMA8, PSMB1, PSMB10, PSMB2, PSMB3, PSMB4, PSMB5, PSMB6, PSMB7, PSMB8, PSMB9, PSMC1, PSMC2, PSMC3, PSMC4, PSMC5, PSMC6, PSMD1, PSMD10, PSMD11, PSMD12, PSMD13, PSMD14, PSMD2, PSMD3, PSMD4, PSMD5, PSMD6, PSMD7, PSMD8, PSMD9, PSME1, PSME2, PSME4, PSMF1, PTTG1, RPS27A, SKP2, UBA52, UBE2C, UBE2D1, UBE2E1 |
| Genes involved in Beta-catenin phosphorylation cascade | APC, AXIN1, CSNK1A1, CTNNB1, FAM123B, FRAT1, FRAT2, PPP2CA, PPP2CB, PPP2R1A, PPP2R1B, PPP2R5A, PPP2R5B, PPP2R5C, PPP2R5D, PPP2R5E |
| Autodegradation of Cdh1 by Cdh1:APC/C | ANAPC1, ANAPC10, ANAPC11, ANAPC2, ANAPC4, ANAPC5, ANAPC7, CDC16, CDC23, CDC26, CDC27, PSMA1, PSMA2, PSMA3, PSMA4, PSMA5, PSMA6, PSMA7, PSMB1, PSMB10, PSMB2, PSMB3, PSMB4, PSMB5, PSMB6, PSMB7, PSMB8, PSMB9, PSMC1, PSMC2, PSMC3, PSMC4, PSMC5, PSMC6, PSMD1, PSMD10, PSMD11, PSMD12, PSMD13, PSMD14, PSMD2, PSMD3, PSMD4, PSMD5, PSMD6, PSMD7, PSMD8, PSMD9, PSME1, PSME2, PSMF1, RPS27A, UBA52, UBE2C, UBE2D1, UBE2E1 |
| Genes involved in M/G1 transition | CDC45, CDC6, CDC7, CDK2, CDT1, DBF4, E2F1, E2F2, E2F3, GMNN, MCM10, MCM2, MCM3, MCM4, MCM5, MCM6, MCM7, MCM8, POLA1, POLA2, POLE, POLE2, PRIM1, PRIM2, PSMA1, PSMA2, PSMA3, PSMA4, PSMA5, PSMA6, PSMA7, PSMA8, PSMB1, PSMB10, PSMB2, PSMB3, PSMB4, PSMB5, PSMB6, PSMB7, PSMB8, PSMB9, PSMC1, PSMC2, PSMC3, PSMC4, PSMC5, PSMC6, PSMD1, PSMD10, PSMD11, PSMD12, PSMD13, PSMD14, PSMD2, PSMD3, PSMD4, PSMD5, PSMD6, PSMD7, PSMD8, PSMD9, PSME1, PSME2, PSME4, PSMF1, RPA1, RPA2, RPA3, RPA4, RPS27A, UBA52 |
| Regulation of the Fanconi anemia pathway | ATM, ATR, FANCD2, RPS27A, UBA52, USP1, ZBTB32 |
| Apoptotic cleavage of cellular proteins | ACIN1, ADD1, APC, BCAP31, BIRC2, BMX, CASP3, CASP6, CASP7, CASP8, CDH1, CTNNB1, DBNL, DSG1, DSG2, DSG3, DSP, FNTA, GAS2, GSN, LMNA, LMNB1, LOC647859, MAPT, MST4, OCLN, PKP1, PLEC, PRKCD, PRKCQ, PTK2, ROCK1, SATB1, SPTAN1, STK24, TJP1, TJP2, VIM |
| Apoptosis | ACIN1, ADD1, AKT1, APAF1, APC, APPL1, ARHGAP10, BAD, BAK1, BAX, BBC3, BCAP31, BCL2, BCL2L1, BCL2L11, BID, BIRC2, BMF, BMX, CASP10, CASP3, CASP6, CASP7, CASP8, CASP9, CDH1, CFLAR, CTNNB1, CYCS, DAPK1, DAPK2, DAPK3, DBNL, DCC, DFFA, DFFB, DIABLO, DNM1L, DSG1, DSG2, DSG3, DSP, DYNLL1, DYNLL2, E2F1, FADD, FAS, FASLG, FNTA, GAS2, GSN, GZMB, H1F0, HIST1H1A, HIST1H1B, HIST1H1C, HIST1H1D, HIST1H1E, HMGB1, HMGB2, KPNA1, KPNB1, LMNA, LMNB1, LOC647859, MAGED1, MAPK8, MAPT, MST4, NMT1, OCLN, PAK2, PKP1, PLEC, PMAIP1, PPP3R1, PRKCD, PRKCQ, PSMA1, PSMA2, PSMA3, PSMA4, PSMA5, PSMA6, PSMA7, PSMA8, PSMB1, PSMB10, PSMB2, PSMB3, PSMB4, PSMB5, PSMB6, PSMB7, PSMB8, PSMB9, PSMC1, PSMC2, PSMC3, PSMC4, PSMC5, PSMC6, PSMD1, PSMD10, PSMD11, PSMD12, PSMD13, PSMD14, PSMD2, PSMD3, PSMD4, PSMD5, PSMD6, PSMD7, PSMD8, PSMD9, PSME1, PSME2, PSME4, PSMF1, PTK2, RIPK1, ROCK1, RPS27A, SATB1, SPTAN1, STK24, TFDP1, TJP1, TJP2, TNF, TNFRSF10B, TNFRSF1A, TNFSF10, TP53, TRADD, TRAF2, UBA52, UNC5A, UNC5B, VIM, XIAP, YWHAB |
| ER-phagosome pathway | B2M, CALR, HLA-A, HLA-B, HLA-C, HLA-F, HLA-G, PDIA3, PSMA1, PSMA2, PSMA3, PSMA4, PSMA5, PSMA6, PSMA7, PSMA8, PSMB1, PSMB10, PSMB2, PSMB3, PSMB4, PSMB5, PSMB6, PSMB7, PSMB8, PSMB9, PSMC1, PSMC2, PSMC3, PSMC4, PSMC5, PSMC6, PSMD1, PSMD10, PSMD11, PSMD12, PSMD13, PSMD14, PSMD2, PSMD3, PSMD4, PSMD5, PSMD6, PSMD7, PSMD8, PSMD9, PSME1, PSME2, PSME4, PSMF1, RPS27A, SEC61A1, SEC61A2, SEC61B, SEC61G, TAP1, TAP2, UBA52 |

Supplementary Table 6. Median and mean (across 200 simulated datasets) values for the area under the receiver operating characteristic curve (AUC) for identifying aberrant individuals using pathway deviation scores calculated with multi-omic *padma*, single-omics *padma*, and a PCA of concatenated data for different simulation settings ($n$ = 30, 50, 100, 250, 500; $p_{aberrant}$ = 1% or 5%). Results are shown for $|d_{genes}|$ = 2 and $|d_{omics}|$ = 1 (i.e. two driver genes, with a single driver omic per gene, for each aberrant individual). Best performers for each combination of simulation parameters are highlighted in yellow.

| *n* | $p_{aberrant}$ | Method | Median AUC | Mean AUC |
|---|---|---|---|---|
| 30 | 1% | *padma* | 0.862 | 0.818 |
| | | *padma* single-omics | 0.845 | 0.811 |
| | | PCA | 0.828 | 0.802 |
| | 5% | *padma* | 0.768 | 0.765 |
| | | *padma* single-omics | 0.679 | 0.698 |
| | | PCA | 0.732 | 0.749 |
| 50 | 1% | *padma* | 0.816 | 0.813 |
| | | *padma* single-omics | 0.837 | 0.815 |
| | | PCA | 0.796 | 0.796 |
| | 5% | *padma* | 0.787 | 0.783 |
| | | *padma* single-omics | 0.670 | 0.682 |
| | | PCA | 0.773 | 0.771 |
| 100 | 1% | *padma* | 0.879 | 0.827 |
| | | *padma* single-omics | 0.859 | 0.826 |
| | | PCA | 0.864 | 0.823 |
| | 5% | *padma* | 0.802 | 0.796 |
| | | *padma* single-omics | 0.647 | 0.659 |
| | | PCA | 0.800 | 0.789 |
| 250 | 1% | *padma* | 0.844 | 0.828 |
| | | *padma* single-omics | 0.671 | 0.691 |
| | | PCA | 0.841 | 0.825 |
| | 5% | *padma* | 0.822 | 0.818 |
| | | *padma* single-omics | 0.642 | 0.641 |
| | | PCA | 0.817 | 0.813 |
| 500 | 1% | *padma* | 0.849 | 0.833 |
| | | *padma* single-omics | 0.655 | 0.658 |
| | | PCA | 0.845 | 0.833 |
| | 5% | *padma* | 0.828 | 0.828 |
| | | *padma* single-omics | 0.653 | 0.652 |
| | | PCA | 0.825 | 0.825 |
